# Development of Learning Objectives for Non-Major Introductory Biology Using a Delphi Method

**DOI:** 10.1101/2024.12.19.629465

**Authors:** Peggy Brickman, Cara Gormally

**Affiliations:** University of Georgia; Gallaudet University

**Keywords:** Learning Objectives, Delphi Method, Non-Science Majors, Socioscientific Issues, Science Competencies

## Abstract

Biology faculty have consensus-based guidelines based on Vision and Change principles about what to teach introductory biology majors. In contrast, faculty have not reached a consensus concerning the learning goals for introductory non-majors courses. Yet, more than 8 out of 10 undergraduates are not science majors. The goal of this study was to develop and evaluate learning objectives for non-majors introductory biology courses. We performed a modified-Delphi study of learning objectives (LOs) for non-majors biology. We engaged a total of 38 biology faculty experts from institutions across the US in three iterative rounds to identify, rate, discuss, and re-rate >300 LOs for non-majors biology courses. Faculty provided feedback to determine whether the LOs are critical for students to learn and if the LOs encompass what students need to learn about this issue, as well as if anything is missing. As a result of expert evaluation, 60.7% of LOs (164) were deemed critical. Experts also suggested 22 additional new LOs.

## Introduction

Educators face a universal conundrum: identifying what is important for students to learn. This is particularly challenging for faculty teaching courses designed to support science literacy rather than providing a foundation for future coursework or professional preparation. Through a large consensus of faculty voices, Vision & Change transformed biology education by establishing goals and themes directing what undergraduate biology majors should learn (AAAS, 2010). This landmark event disrupted the status quo of focusing solely on content coverage in science teaching in the United States of America. In contrast, however, faculty have not reached a consensus concerning the learning goals for non-majors courses. Yet, more than 8 out of 10 college students are not science majors (Statistics, 2019). As part of their general education requirements, most undergraduates in the U.S. take a natural science course. Most will likely never take another science course (Statistics, 2019). Biology faculty tend to overlook this huge population of students in lieu of preparing STEM majors (Gormally & Heil, 2022). When we recognize this reality, determining what is important for non-majors to learn is of utmost urgency to support a science literate public.

Making science learning useful is incredibly important for non-majors as we face escalating climate concerns, health disparities, and declining trust in science (Kennedy & Tyson, 2023). Our future hinges on these future business owners, educators, and politicians who will navigate these looming socio-scientific crises. When surveyed about how science education can be improved for Vision and Change, undergraduates requested the presence of more topic-based or concept-oriented courses, especially for non-majors courses (Brewer & Smith, 2011). Research suggests strategies to make science learning useful, specifically, prioritizing socio-scientific issues, highlighting communal opportunities in science that impact students’ communities, and providing students with opportunities to practice skills to engage with science beyond the classroom (Gormally & Heil, 2022; Stephens et al., 2017) . However, from analyzing syllabi, 48% of non-majors biology courses (N=78) focus solely on science content, with little attention to socio-scientific issues (Heil et al., 2024). Consequently, what non-majors learn may not be directly relevant to their everyday lives beyond the biology classroom. Notably, as the Vision & Change report cites, “regardless of their ultimate career paths, all students will need these very basic skills [of using evidence and logic to reach sound conclusions] to participate as citizens and thrive in the modern world.”

Learning objectives (hereafter referred to as LOs) are statements of what students should know and be able to do at the end of a specific class session (Orr et al., 2022). LOs are the foundation of backward design. In a well-designed course, learning objectives, opportunities for students to practice, and assessments should all be aligned (Wiggins & McTighe, 2011).

Comprehensive course design–aligning active learning exercises with learning objectives and assessments–offers faculty a way forward through a clear articulation of what is important for students to learn.

Critically, these learning objectives must provide integrated opportunities to practice these skills with conceptual learning.

Most instructors do not design courses around LOs but instead create courses that focus on topics and content coverage (Heil et al., 2024). Additionally, lesson-level learning outcomes from non-majors biology courses rarely were aligned with a competency skill, were more likely to be lower-level (i.e., remember and understand), and were rarely tied to a socio-scientific issue (Heil et al., 2023). When instructors do use LOs, they often use learning objectives that are mandated—and these can vary in quality and in their level of cognitive challenge (Heil et al., 2023). Learning objectives are often used for U.S. articulation agreements to dictate transfer of credits between 2- and 4-year colleges, but these can vary in quality (Lennon, 2018). Notably, non-major courses fulfill general education requirements. This means that learning objectives can provide clarity and oversight to be sure these goals are met in the course, especially as these courses may be taught by multiple instructors, in multiple sections.

Our research project was designed to facilitate an increased emphasis on the directive set forth by Vision & Change to “ensure that undergraduate biology courses are active, outcome-oriented, inquiry-driven, and relevant” with a “focus on conceptual understanding, not just on covering voluminous content.” Additionally, we aimed to integrate competencies as articulated in BioSkills Guide with socio-scientific issues to make biology learning useful and relevant to students’ real world lives (Brewer & Smith, 2011; Clemmons et al., 2020; Feinstein et al., 2013). In a recent analysis of instructors’ learning objectives in introductory biology courses for non-majors, competency skills were present in only 17.7% of instructors’ LOs and 7% of the textbook LOs (Heil et al., 2023). We aimed to specifically solicit input from faculty with expertise in teaching non-majors who have considered the importance of these aspects of Vision & Change and teach issues-based courses. To write the LOs, we followed best practices as described by Orr et al (2022). We aimed to develop LOs appropriate for faculty to use to design a single class session.

### Methodological Rationale

Since our research question required faculty experts, we used the Delphi Method. The Delphi method is an iterative process involving multiple rounds of data collection to elicit expert input and explore areas of consensus. Typically, Delphi participants anonymously provide answers to a survey. Then, survey responses are analyzed using descriptive statistics. Responses are shared with the Delphi participants, who then answer a second set of questions. This process is repeated until either consensus is reached or a predetermined number of rounds have passed, which mitigates some problems associated with expert panels (Ven & Delbecq, 1974).

We approached our research question with an online modified-Delphi study. First, we solicited participants’ feedback via a Qualtrics survey. Then, we used Google docs to share results with participants to evaluate and reach an agreement about the LOs. This approach combines a traditional Delphi structure with a round of online asynchronous discussion among participants. This asynchronous approach uses our expert participants’ time efficiently (Bowles et al., 2003), and encourages more individual contribution as well as limits counterproductive behaviors such as groupthink or influence of participant status (Dubrovsky et al., 1991; Murphy et al., 1998; Pagliari et al., 2001).

Typically, two to three rounds of participant feedback suffice to achieve consensus if it exists (Woudenberg, 1991). More than three rounds are associated with increased participant burden and boredom, as well as significant declines in participation rates (Keeney et al., 2001). There is no agreement about optimal size for Delphi studies (Murry Jr & Hammons, 1995; Rowe, 2001). Large numbers of Delphi participants may be difficult to coordinate (Murphy et al., 1998). Small numbers may result in lower quality of discussion (Vonderwell, 2003). Prior work suggests that approximately 40 participants is a good number for a productive online discussion (Khodyakov et al., 2011). Thus, our study aimed to include approximately 40 participants.

## Methods

### Overview of the process

Our main study goal was to develop a set of LOs that align with broader course goals directed by Vision & Change. Specifically, we aimed to develop enough LOs to create a large selection of common content through the lens of socio-scientific issues. As a result, these LOs are designed to provide instructors latitude to customize their course content for their program and student population. Instructors can select specific socio-scientific issues and the associated LOs that are of most interest for their courses.

The process was divided into two broad phases: a development phase and an evaluation phase (Figure 1) similar to Hennesey & Freeman (2024). During the development phase, candidate LOs went through multiple rounds of evaluation and revision. In total, there were three different groups of researcher-instructors that participated in the development phase and another Delphi group that participated in 2 rounds of review during the evaluation phase. Each group involved different teams of evaluators, all of whom shared instructional expertise in life sciences content and experience with biology education research:

- Group One was composed of this manuscript’s authors who drafted an initial set of LOs designed to align with issues identified in an analysis of syllabi collected nationally (Heil et al., 2024) which were also compared to a set of LOs developed for introductory biology courses designed for biology majors (Hennessey & Freeman, 2024).
- Group Two was composed of 5 educators who teach non-majors Biology and regularly advise the BioInteractive program at the Howard Hughes Medical Institute (HHMI) on curriculum development.
- Group Three consisted of two experts with extensive experience in writing LOs and in assessment design.

**Figure 1:**
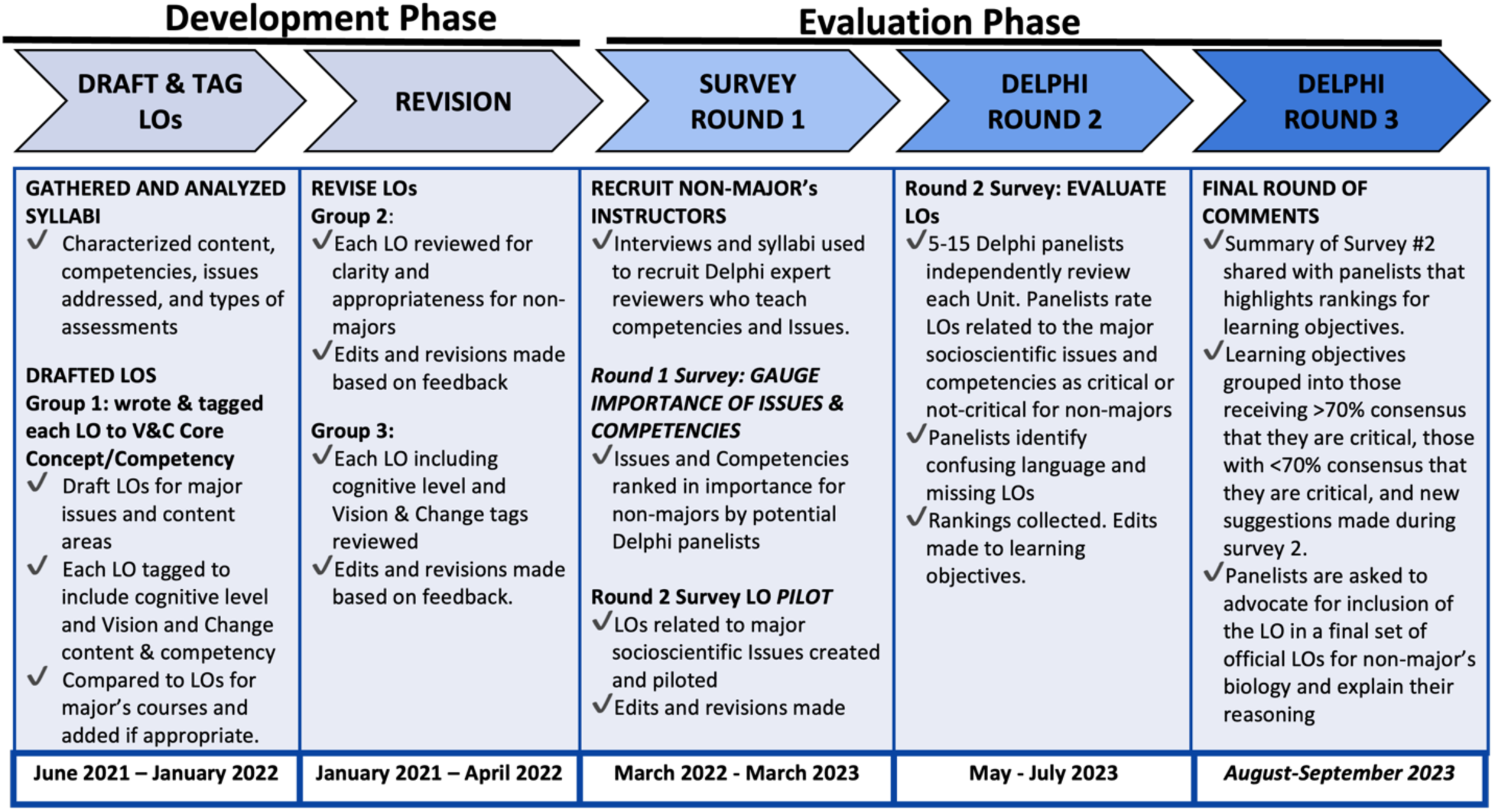
Development and evaluation phases of learning objectives for introductory biology for non-science majors.

### Development Phase

We drafted LOs for six units based on the organization and content presented as major areas defined in our analysis of syllabi (Heil et al., 2024). The units included: Biochemistry, Cell Biology, Genetics, Evolution, Animal Physiology, and Ecology. Individual LOs were drafted to adequately describe both topics and socio-scientific issues commonly taught in issues-based courses from our national survey (Supplemental Materials, Table 1). Each author independently rated each candidate LO as representing either lower order cognitive skill (LOCS) or higher order cognitive skills (HOCS) based on Bloom’s taxonomy for learning (Anderson & Krathwohl, 2001), paired LOCS and HOCS LOs on the same concept whenever possible, and aligned each LO to one or more of the concepts and competencies articulated in the final Vision and Change Report (Brewer & Smith, 2011) as well as one or more statements in the BioCore Guide and BioSkills Guide (Brownell et al., 2014; Clemmons et al., 2020). Finally, the authors met to discuss and reach agreement in each LOCS versus HOCS designation, proposed LOCS-HOCS pairing, and Vision and Change, BioCore Guide, and BioSkills Guide tags.

**Table 1.** Learning Objectives Reviewed Compared to Final Recommended Set.

| **Phase** | **UNIT 1**<br>**Cell**<br>**Biology** | **UNIT 2**<br>**Biochemi**<br>**stry** | **UNIT 4**<br>**Evolution** | **UNIT 3**<br>**Genetics** | **UNIT 5**<br>**Animal**<br>**Physiology** | **UNIT 6**<br>**Ecology** | **Total** |
| --- | --- | --- | --- | --- | --- | --- | --- |
| **Group 1** | 25 LOCS<br>11 HOCS | 23 LOCS<br>14 HOCS | 21 LOCS<br>7 HOCS | 30 LOCS<br>27 HOCS | 47 LOCS<br>23 HOCS | 55 LOCS<br>35 HOCS | **318** |
| **Group 2** | 6 LOCS<br>5 HOCS | 17 LOCS<br>10 HOCS | 8 LOCS<br>5 HOCS | 33 LOCS<br>28 HOCS | 40 LOCS<br>18 HOCS | 43 LOCS<br>39 HOCS | **255** |
| **Group 3** | 11 LOCS<br>7 HOCS | 12 LOCS<br>8 HOCS | 12 LOCS<br>10 HOCS | 23 LOCS<br>25 HOCS | 43 LOCS<br>24 HOCS | 46 LOCS<br>51 HOCS | **270** |
| **Final**<br>**LOs** | 4 LOCS<br>8 HOCS | 9 LOCS<br>7 HOCS | 4 LOCS<br>1 HOCS | 17 LOCS<br>15 HOCS | 31 LOCS<br>13 HOCS | 26 LOCS<br>29 HOCS | **164** |

The candidate LOs that emerged from this initial work (N=318) underwent two rounds of review (Groups 2 & 3) to determine if the reviewers felt that the LO was clear, was useful for student learning about this issue, and was at the appropriate difficulty level for non-majors taking an introductory course (Table 1). Codes were developed to characterize comments left by Group 2 and Group 3 and all comments were coded to agreement by both authors A full list of all codes can be found in the Supplemental Materials (Table 2). The most frequent comments provided by Group 2 related to: (1) the LO being more appropriate for majors (51%); (2) the LO not being useful to students learning about this issue (16%); and (3) rewording for clarity or simplification (13%). The most frequent specific comments left by Group 3 related to: (1) condensing LOs to remove redundancy (37%); (2) changing verbs or clarifying meaning (36%); and (3) confusion about specifics about the LO or its Bloom’s level (24%). After each round of review, the two authors discussed feedback until reaching agreement for each suggested revision. Many revisions involved condensing or removing inappropriate LOs or moving LOs specific to one issue into another unit. In total, each LO went through two rounds of revision during the study’s “Draft & Tag LOs & Revision” step (Figure 1). Changes were made as indicated (Table 1).

**Table 2.** Institutions Represented by Delphi Faculty Expert Participants.

|  | Development Phase | Delphi Evaluation Phase |  |  |  |
| --- | --- | --- | --- | --- | --- |
|  | Issues & Competency Survey | Group 2 | Round 1 | Round 2 | Round 3 |
| **Total Participants** | **20** | **5** | **20** | **26** | **13** |
| **Institution Type** |  |  |  |  |  |
| Doctoral Universities | 55% | 20% | 55% | 58% | 62% |
| Very high research activity | 8 | 1 | 11 | 9 | 4 |
| High research activity | 3 |  | 8 | 5 | 3 |
| Doctoral/Professional University |  |  | 3 | 1 | 1 |
| Master's Colleges and Universities | 15% | 20% | 20% | 12% | 0% |
| Larger Programs | 1 | 1 | 2 | 2 |  |
| Medium Programs |  |  |  |  |  |
| Smaller Programs | 1 |  | 1 | 1 |  |
| Baccalaureate Colleges | 15% | 20% | 15% | 23% | 15% |
| Arts and Sciences Focus | 3 | 1 | 3 | 6 | 2 |
| Diverse Fields |  |  |  |  |  |
| Two-year Institutions | 15% | 40% | 15% | 12% | 23% |
| Community Colleges | 3 | 2 | 3 | 3 | 3 |
| Health Professions |  |  |  |  |  |
| Minority Serving | 0% | 60% | 0% | 12% | 23% |
|  | 0 | 3 | 0 | 3 | 3 |

### Evaluation Phase

#### Recruitment for the Delphi Study

Once the development phase was complete, the LOs went through an evaluation process based on leveraging the expertise of an identified group of experienced instructors who teach introductory biology for non-majors with an issues-based approach (Figure 2). Unlike Hennessey and Freeman (2024), we used a Delphi approach because our previous examination of a national sample of U.S. syllabi indicated that most faculty teach non-majors using a content-based approach that does not align with recommendations made by Vision & Change to include “active, outcome-oriented, inquiry-driven, and relevant” with a “focus on conceptual understanding, not just on covering voluminous content.” Faculty participants were asked to assess and rate each LO as critical or not critical for introductory biology for non-majors courses, while recognizing that each faculty member would also contribute their own LOs to customize their course to their institution and student population.

**Figure 2.**
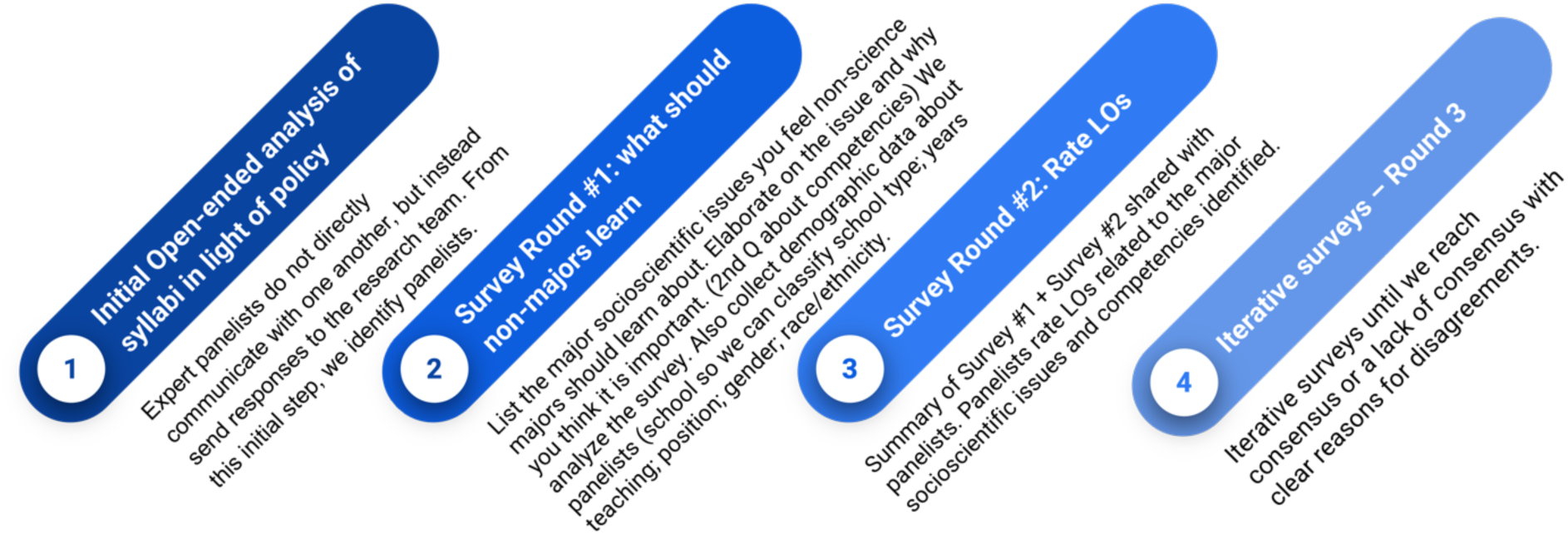
Overview of the Delphi Method.

We contacted 244 instructors of non-majors introductory biology in the U.S. in our attempt to identify our panel of experts (Table 2). Potential participants were identified from numerous sources, including: a prior study of syllabi (Heil et al., 2024); BioInteractive Higher Ed Newsletter Subscribers; the Partnership for Undergraduate Life Sciences Education community; the Society for the Advancement of Biology Education Research; a list of biology faculty from HBCU and tribal colleges; and the American Society for Cell Biology Education Group. Additionally, we attempted to identify potential participants via a snowballing method where we asked instructors who were involved in other NSF-funded projects involving non-majors biology (Improvement of General Education Life Science courses (IGELS) and ORACLE) to recommend colleagues that taught non-majors. Interested individuals were also encouraged to share the survey. We began recruiting participants in May of 2022 and continued to attempt to recruit participants until September 2023. To participate in the survey, respondents were first asked to confirm that they had taught an introductory biology course for non-majors. This study was deemed exempt from Institutional Review Board (IRB) review by the University of Georgia (*PROJECT00003761)* due to the nature of the research, which involved collecting data using anonymous surveys.

### Delphi Study: Surveys

We followed best practices in survey design (Stern et al., 2014) and the principles of social design theory during the Survey Design step highlighted in Figure 1. After developing a preliminary design for the survey in the Qualtrics platform, we engaged Group Three in providing feedback on survey design via written comments and cognitive interviews. We revised the general survey format based on these recommendations and in response to information from think-aloud interviews we conducted with two other colleagues as they took the initial version of the survey. As a final test of the survey, we had one member of Group Three review and provide feedback on each block of the draft survey and comment on both the survey design and the LOs. We revised both the survey and the LOs based on this feedback, resulting in a final format and structure for the survey instrument.

### Delphi Survey Completion

We invited our panel of expert Delphi instructors to participate in the final evaluation steps. instructors were invited to complete a series of surveys (Supplemental Materials, Surveys) for the purpose of reaching a consensus regarding which issues and competencies were most important and to validate a final set of learning objectives for non-science majors. Surveys were piloted (n=2) using a “think aloud,” to identify confusing questions and modify them accordingly. Demographic questions were also included in the first survey, both for tracking purposes, and eventual data correlation purposes. The round 1 survey asked a variety of questions related to the importance of issues, competencies, and learning objectives in non-majors introductory biology and was used to check the balance of issues with content coverage. The round 2 survey offered respondents the opportunity to self-select a unit to evaluate based on their current or prior teaching experience and expertise. Each unit was composed of 1-3 blocks of LOs, based on the total number assigned to that unit. Respondents then evaluated each LO as “critical” or “not critical” to the course they teach. They were also invited to share any feedback about the content or wording of the LOs they evaluated. Respondents were then given an opportunity to return to the survey start and evaluate an additional block of LOs. Respondents did not see any of the data on alignment with Bloom’s level, Vision and Change Core Concepts and Competencies, BioCore Guide statements, or BioSkills Guide statements. Before closing out, the survey also collected institutional and demographic data from each respondent. Between 5-15 panelists reviewed each section. A complete copy of round 1 and 2 surveys is available in the Supplemental Materials.

The final step (round 3) in the evaluation process was for Delphi panelists to be presented with a summary of the results of the round 2 survey that highlights the rankings of critical or not-critical provided for the entire set of learning objectives. Learning objectives grouped into those receiving >70% consensus that they are critical, those with <70% consensus that they are critical, and new suggestions made during Round 2 survey. Panelists are asked to advocate for inclusion of the LO in a final set of official LOs for non-majors biology and explain their reasoning.

## Results

Using a modified Delphi approach, in total 37 unique faculty provided data on their institution and demography (Table 2). Some faculty participated in multiple rounds of surveys. Institution type corresponds to U.S. Carnegie classifications (Shulman, 2001). Percentages may not equal 100 due to rounding. In total, 19% of respondents came from two-year institutions, 11% from baccalaureate institutions, 14% from master’s colleges and universities, and 57% from doctoral universities. If we assume that each of the 2617 total institutions in the 2023 Carnegie Classifications offered a biology course, our data underrepresents two-year, baccalaureate and masters institutions (44% for two year institutions, 22% for baccalaureate, and 25% for masters in the national sample) and over represents doctoral institutions (57% compared to 15% nationally) (Science & Statistics, 2023).

### Delphi Round 1 Survey

A total of 20 faculty experts completed the first 20-minute survey (Qualtrics) that asked faculty to rank both socio-scientific issues (Supplemental Materials, Table 3) and Vision & Change Competencies (Supplemental Materials, Table 4) based on whether these issues and competencies were critical for non-major biology students to learn. At least 65% of faculty ranked the following topics as critical for non-majors to learn: antibiotic resistance and microbiomes; how race is and is not biological; apocalyptic pandemics– immunity; and climate change, C-cycles, biofuels (Supplemental Materials, Table 3).

**Table 3:**
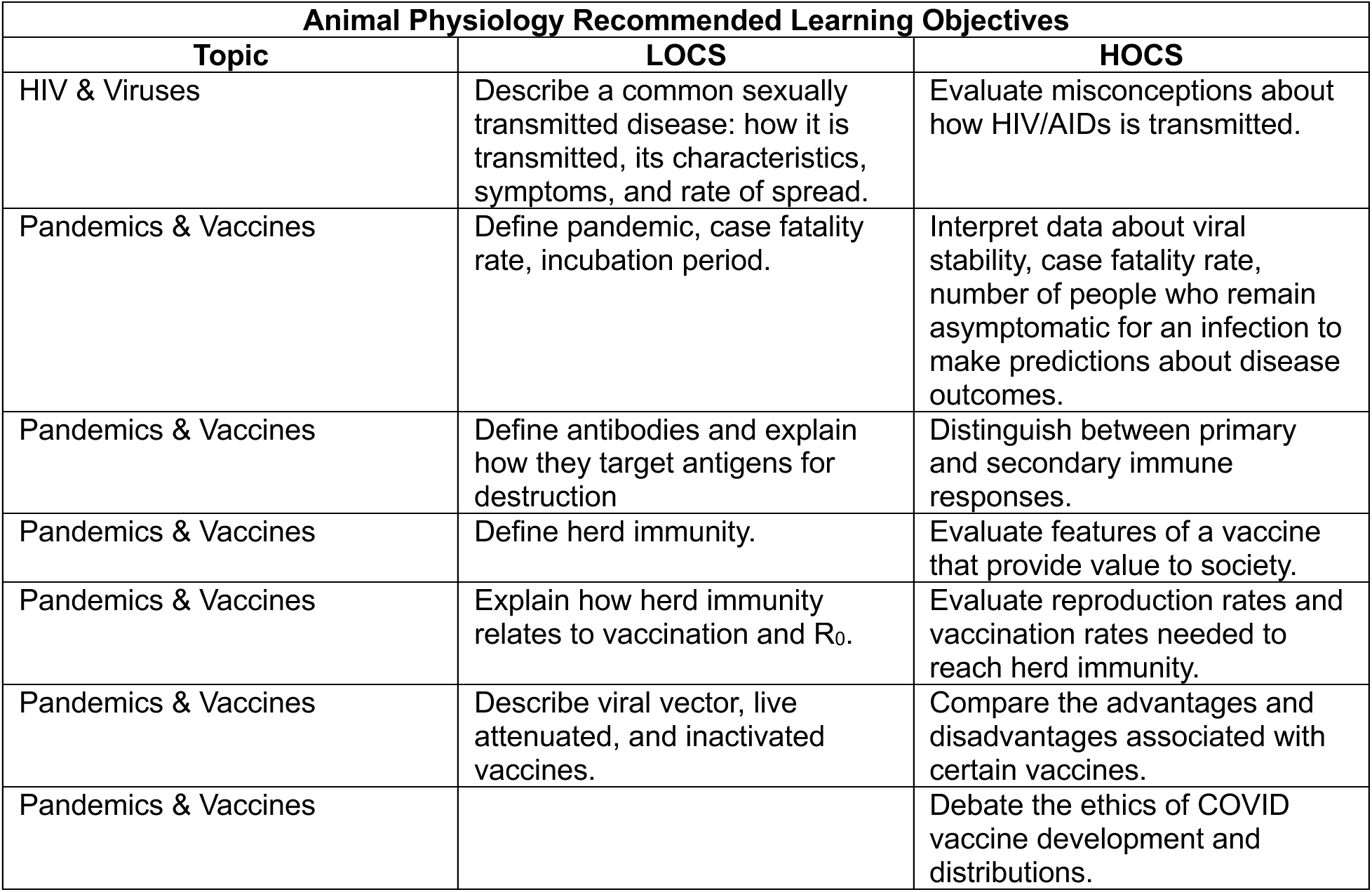
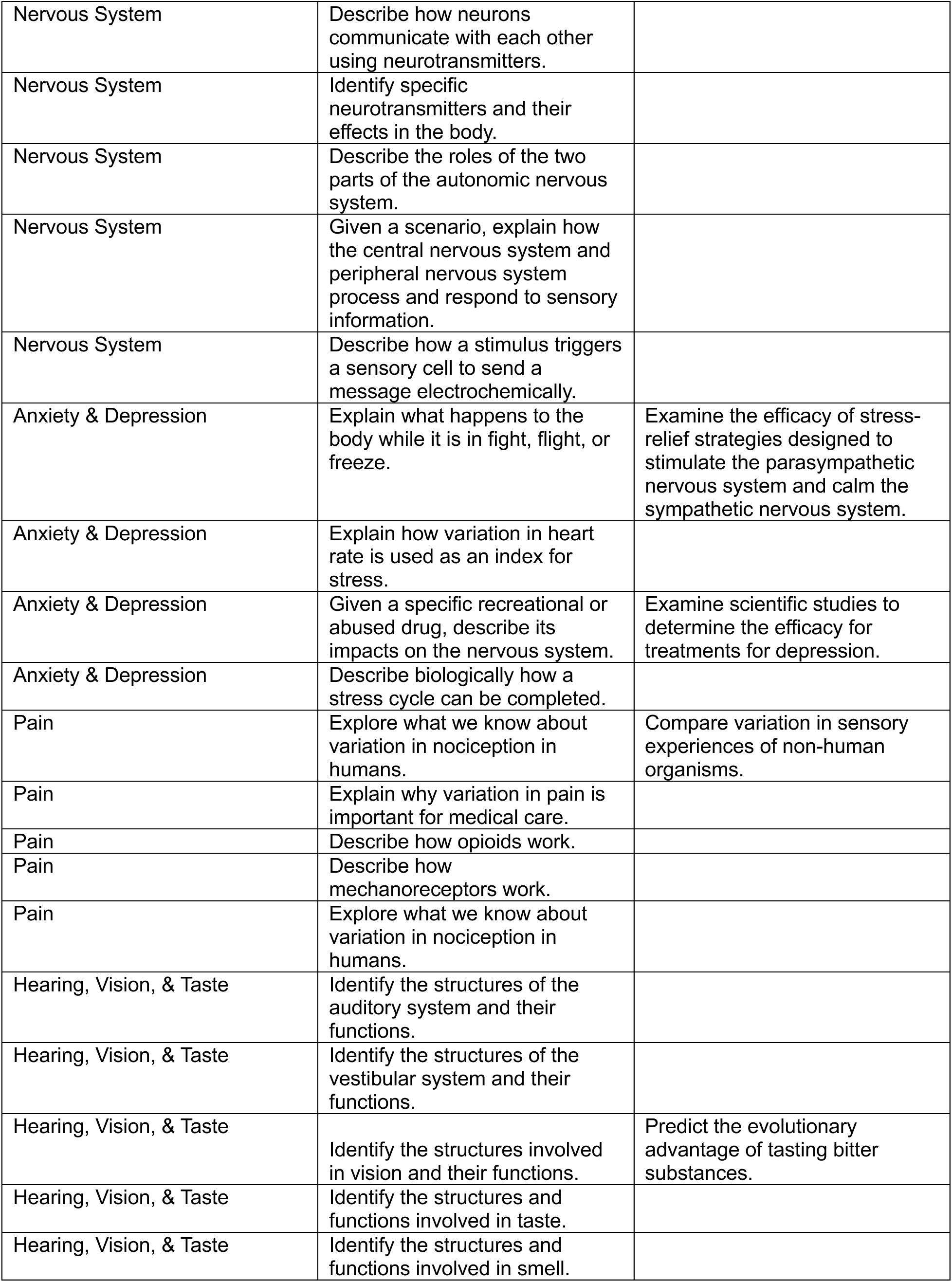
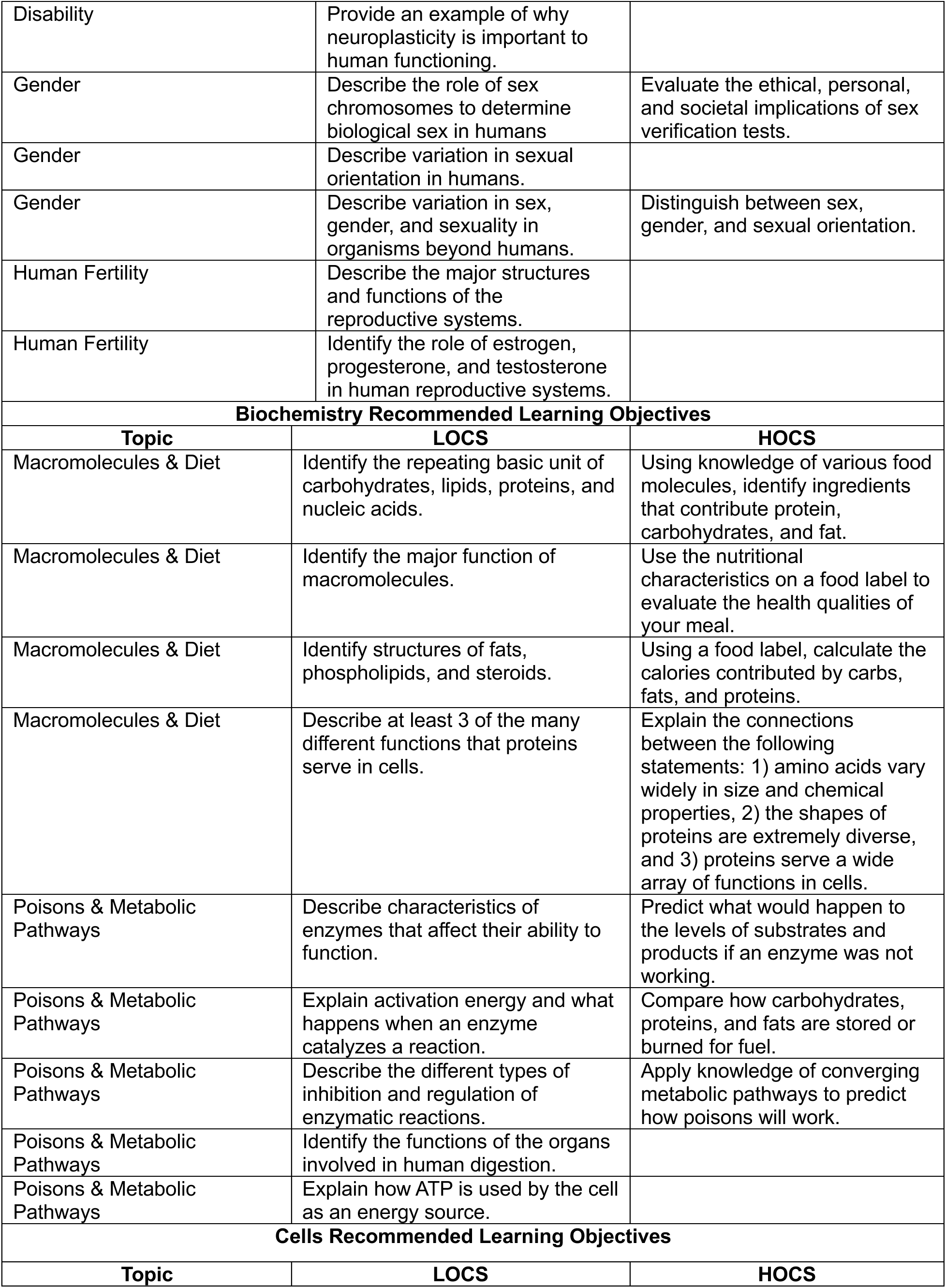
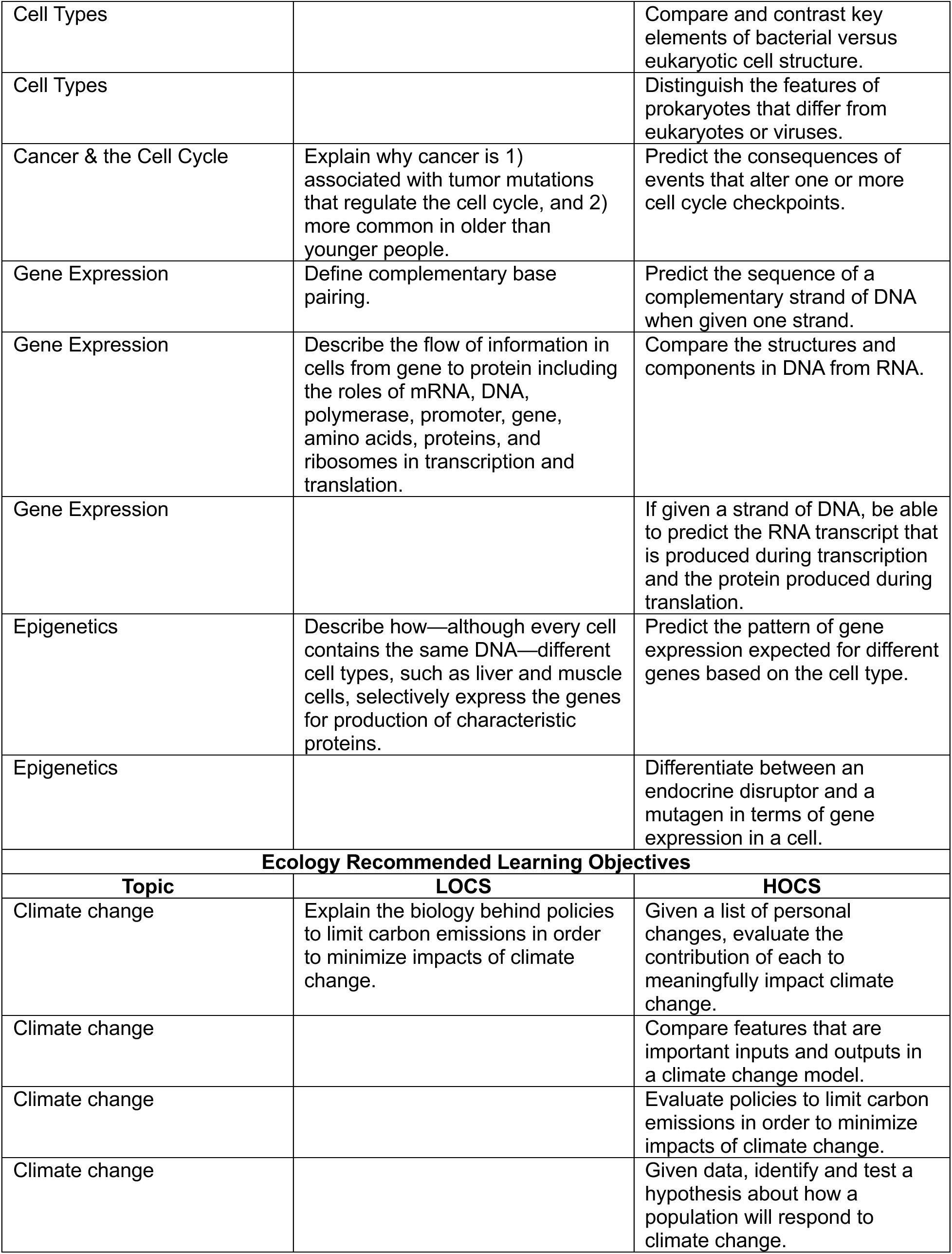
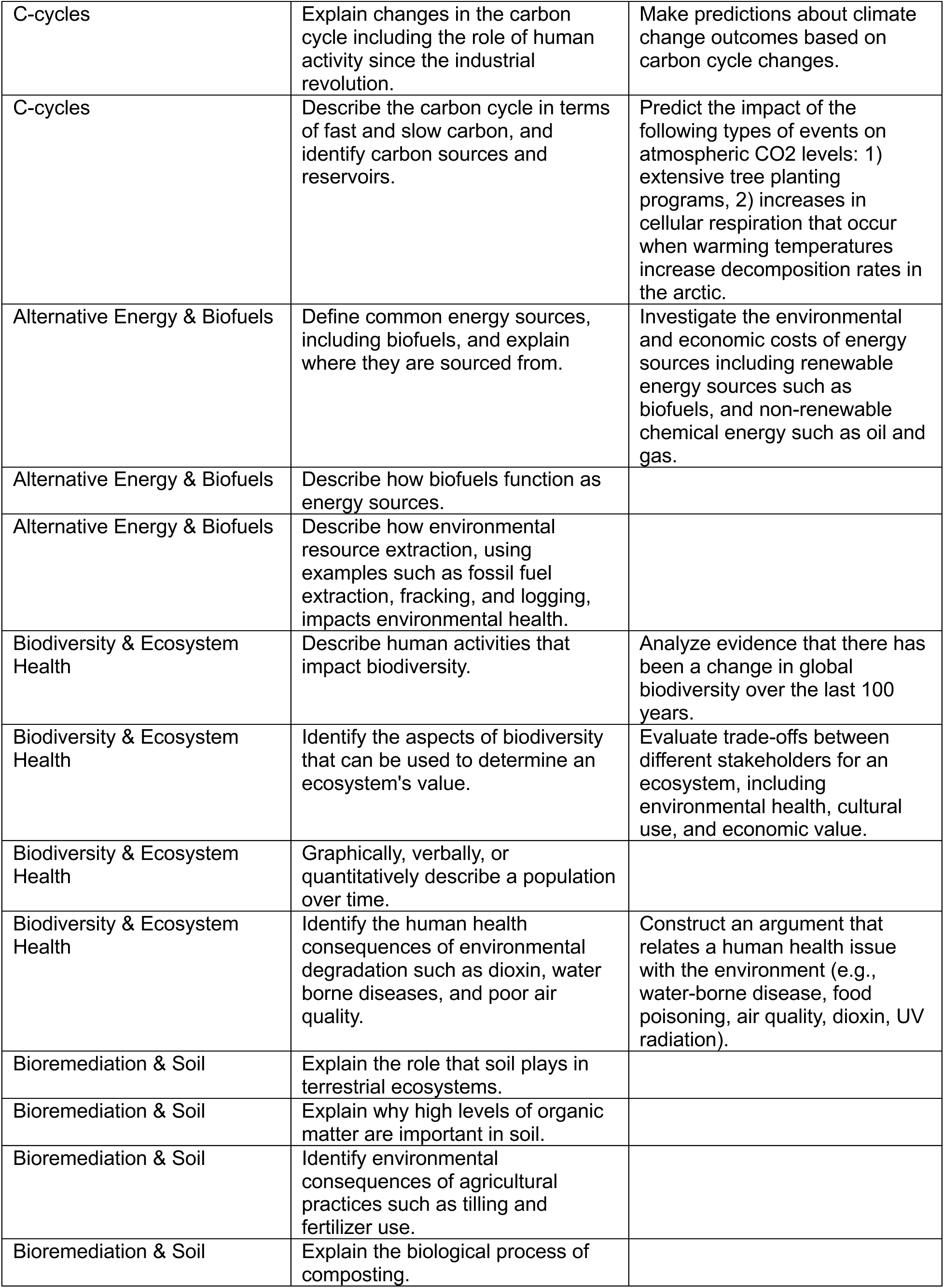
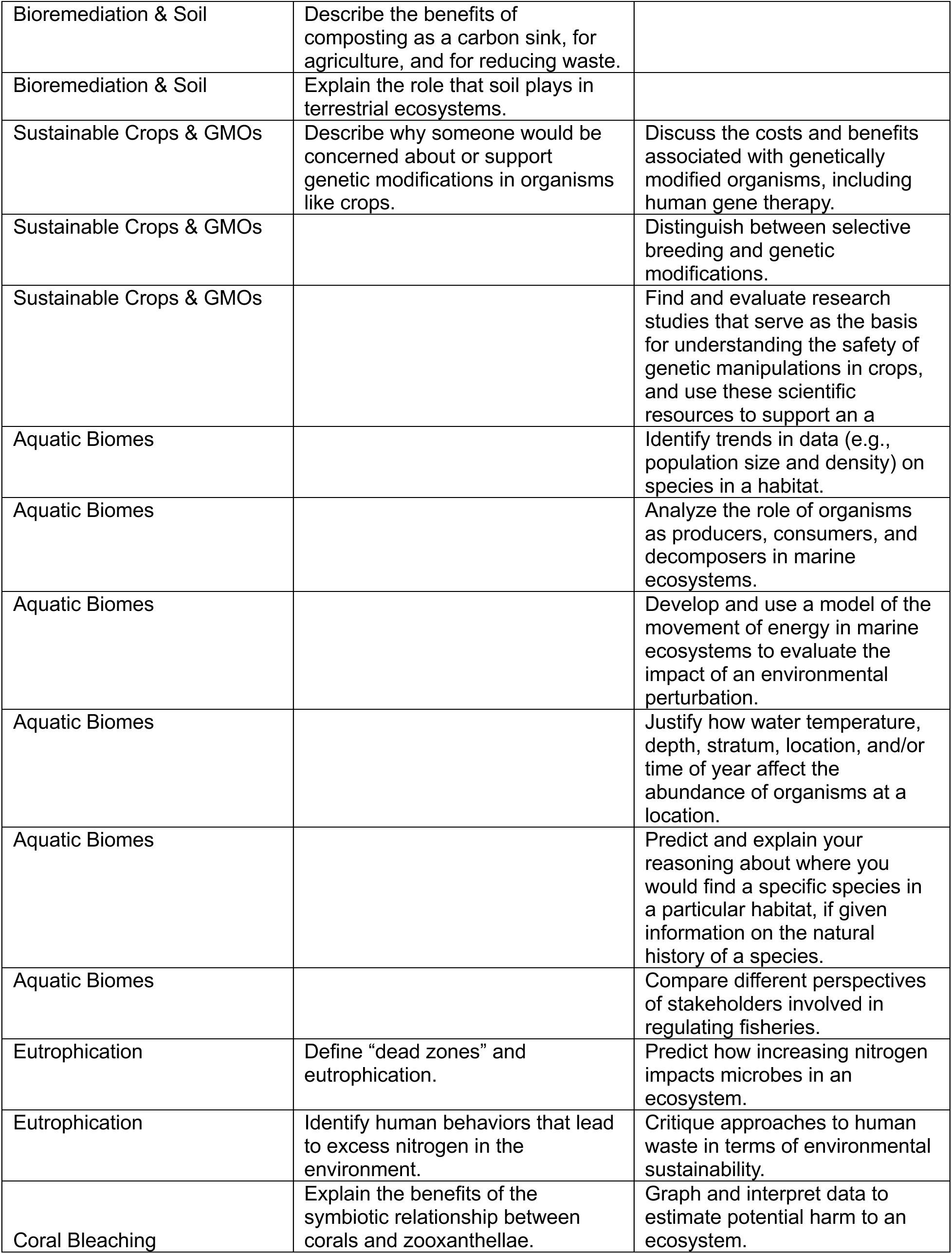
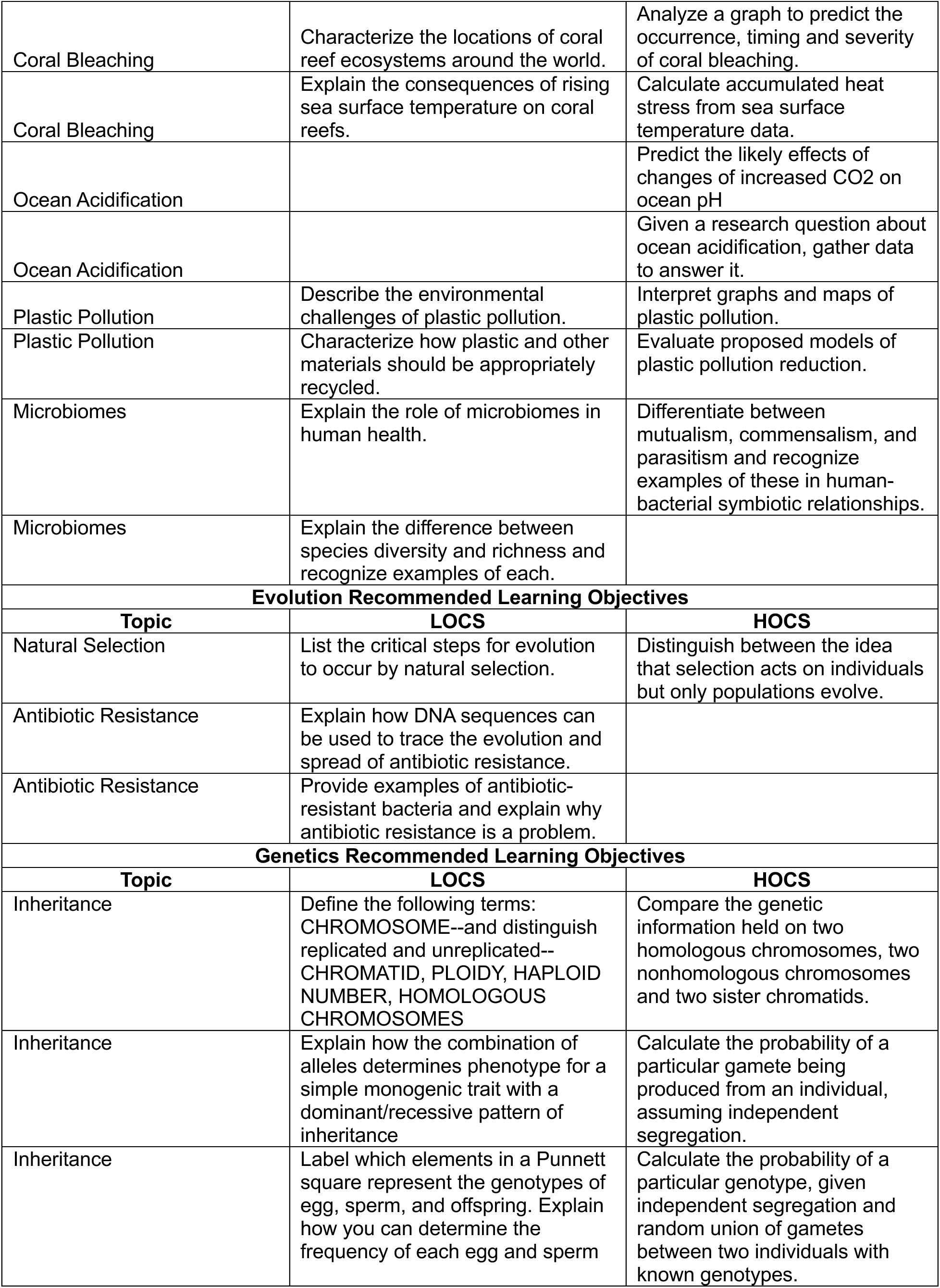
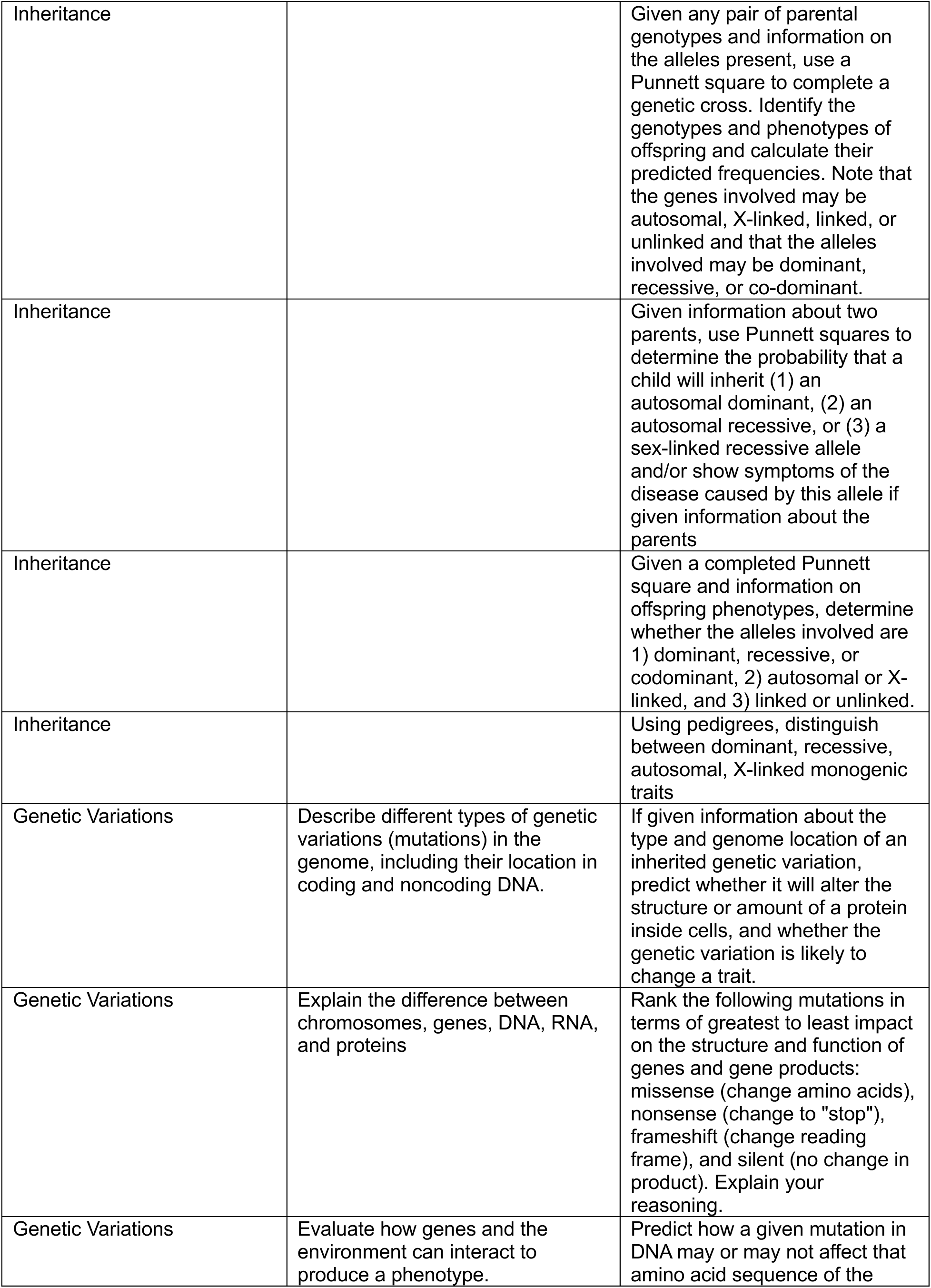
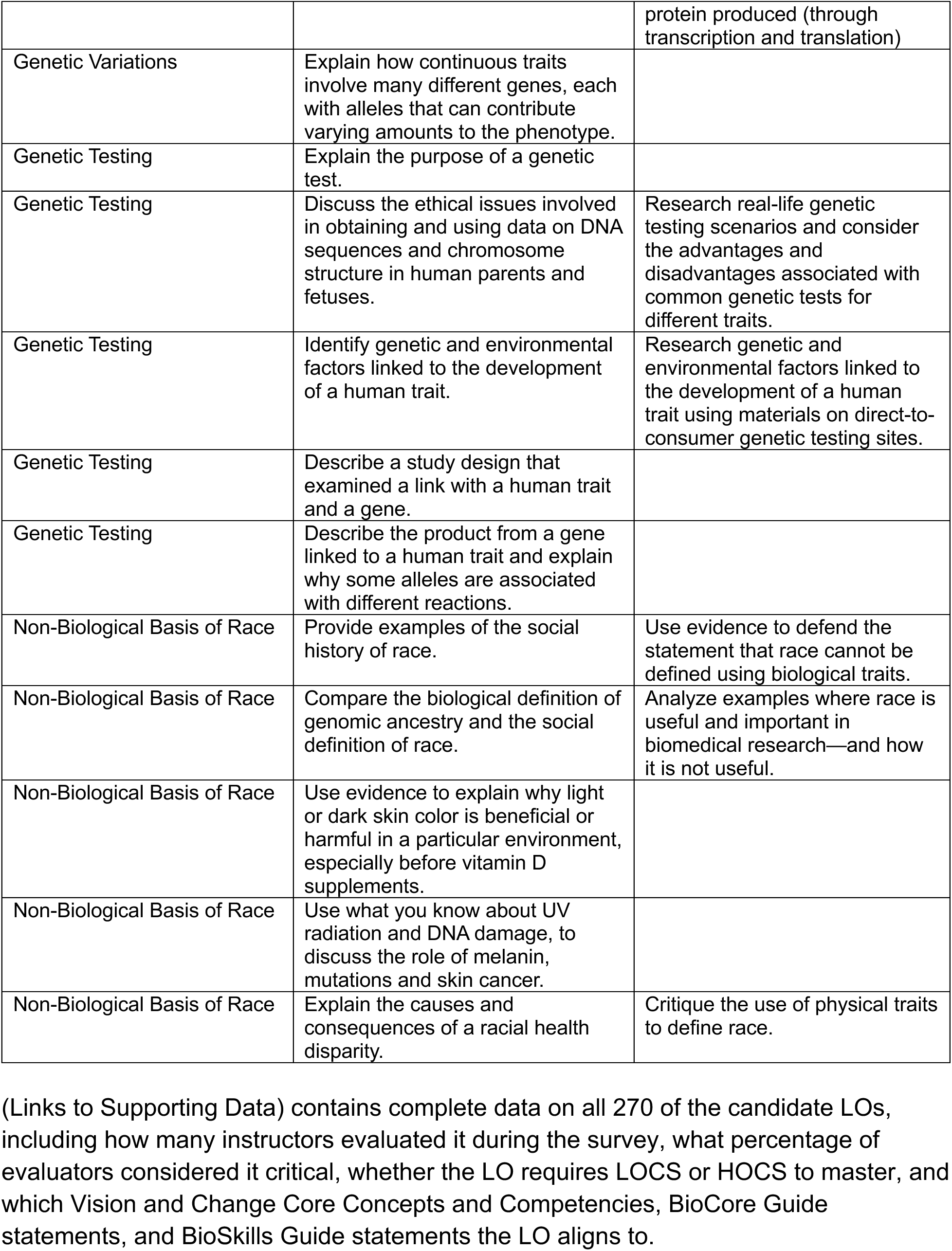
Complete List of Learning Objectives for Introductory Biology for Non-Majors.

| **Animal Physiology Recommended Learning Objectives** |  |  |
| --- | --- | --- |
| **Topic** | **LOCS** | **HOCS** |
| HIV & Viruses | Describe a common sexually transmitted disease: how it is transmitted, its characteristics, symptoms, and rate of spread. | Evaluate misconceptions about how HIV/AIDs is transmitted. |
| Pandemics & Vaccines | Define pandemic, case fatality rate, incubation period. | Interpret data about viral stability, case fatality rate, number of people who remain asymptomatic for an infection to make predictions about disease outcomes. |
| Pandemics & Vaccines | Define antibodies and explain how they target antigens for destruction | Distinguish between primary and secondary immune responses. |
| Pandemics & Vaccines | Define herd immunity. | Evaluate features of a vaccine that provide value to society. |
| Pandemics & Vaccines | Explain how herd immunity relates to vaccination and $R_0$. | Evaluate reproduction rates and vaccination rates needed to reach herd immunity. |
| Pandemics & Vaccines | Describe viral vector, live attenuated, and inactivated vaccines. | Compare the advantages and disadvantages associated with certain vaccines. |
| Pandemics & Vaccines |  | Debate the ethics of COVID vaccine development and distributions. |
| Nervous System | Describe how neurons communicate with each other using neurotransmitters. |  |
| Nervous System | Identify specific neurotransmitters and their effects in the body. |  |
| Nervous System | Describe the roles of the two parts of the autonomic nervous system. |  |
| Nervous System | Given a scenario, explain how the central nervous system and peripheral nervous system process and respond to sensory information. |  |
| Nervous System | Describe how a stimulus triggers a sensory cell to send a message electrochemically. |  |
| Anxiety & Depression | Explain what happens to the body while it is in fight, flight, or freeze. | Examine the efficacy of stress-relief strategies designed to stimulate the parasympathetic nervous system and calm the sympathetic nervous system. |
| Anxiety & Depression | Explain how variation in heart rate is used as an index for stress. |  |
| Anxiety & Depression | Given a specific recreational or abused drug, describe its impacts on the nervous system. | Examine scientific studies to determine the efficacy for treatments for depression. |
| Anxiety & Depression | Describe biologically how a stress cycle can be completed. |  |
| Pain | Explore what we know about variation in nociception in humans. | Compare variation in sensory experiences of non-human organisms. |
| Pain | Explain why variation in pain is important for medical care. |  |
| Pain | Describe how opioids work. |  |
| Pain | Describe how mechanoreceptors work. |  |
| Pain | Explore what we know about variation in nociception in humans. |  |
| Hearing, Vision, & Taste | Identify the structures of the auditory system and their functions. |  |
| Hearing, Vision, & Taste | Identify the structures of the vestibular system and their functions. |  |
| Hearing, Vision, & Taste | Identify the structures involved in vision and their functions. | Predict the evolutionary advantage of tasting bitter substances. |
| Hearing, Vision, & Taste | Identify the structures and functions involved in taste. |  |
| Hearing, Vision, & Taste | Identify the structures and functions involved in smell. |  |
| Disability | Provide an example of why neuroplasticity is important to human functioning. |  |
| Gender | Describe the role of sex chromosomes to determine biological sex in humans | Evaluate the ethical, personal, and societal implications of sex verification tests. |
| Gender | Describe variation in sexual orientation in humans. |  |
| Gender | Describe variation in sex, gender, and sexuality in organisms beyond humans. | Distinguish between sex, gender, and sexual orientation. |
| Human Fertility | Describe the major structures and functions of the reproductive systems. |  |
| Human Fertility | Identify the role of estrogen, progesterone, and testosterone in human reproductive systems. |  |
| **Biochemistry Recommended Learning Objectives** |  |  |
| **Topic** | **LOCS** | **HOCS** |
| Macromolecules & Diet | Identify the repeating basic unit of carbohydrates, lipids, proteins, and nucleic acids. | Using knowledge of various food molecules, identify ingredients that contribute protein, carbohydrates, and fat. |
| Macromolecules & Diet | Identify the major function of macromolecules. | Use the nutritional characteristics on a food label to evaluate the health qualities of your meal. |
| Macromolecules & Diet | Identify structures of fats, phospholipids, and steroids. | Using a food label, calculate the calories contributed by carbs, fats, and proteins. |
| Macromolecules & Diet | Describe at least 3 of the many different functions that proteins serve in cells. | Explain the connections between the following statements: 1) amino acids vary widely in size and chemical properties, 2) the shapes of proteins are extremely diverse, and 3) proteins serve a wide array of functions in cells. |
| Poisons & Metabolic Pathways | Describe characteristics of enzymes that affect their ability to function. | Predict what would happen to the levels of substrates and products if an enzyme was not working. |
| Poisons & Metabolic Pathways | Explain activation energy and what happens when an enzyme catalyzes a reaction. | Compare how carbohydrates, proteins, and fats are stored or burned for fuel. |
| Poisons & Metabolic Pathways | Describe the different types of inhibition and regulation of enzymatic reactions. | Apply knowledge of converging metabolic pathways to predict how poisons will work. |
| Poisons & Metabolic Pathways | Identify the functions of the organs involved in human digestion. |  |
| Poisons & Metabolic Pathways | Explain how ATP is used by the cell as an energy source. |  |
| **Cells Recommended Learning Objectives** |  |  |
| **Topic** | **LOCS** | **HOCS** |
| Cell Types |  | Compare and contrast key elements of bacterial versus eukaryotic cell structure. |
| Cell Types |  | Distinguish the features of prokaryotes that differ from eukaryotes or viruses. |
| Cancer & the Cell Cycle | Explain why cancer is 1) associated with tumor mutations that regulate the cell cycle, and 2) more common in older than younger people. | Predict the consequences of events that alter one or more cell cycle checkpoints. |
| Gene Expression | Define complementary base pairing. | Predict the sequence of a complementary strand of DNA when given one strand. |
| Gene Expression | Describe the flow of information in cells from gene to protein including the roles of mRNA, DNA, polymerase, promoter, gene, amino acids, proteins, and ribosomes in transcription and translation. | Compare the structures and components in DNA from RNA. |
| Gene Expression |  | If given a strand of DNA, be able to predict the RNA transcript that is produced during transcription and the protein produced during translation. |
| Epigenetics | Describe how—although every cell contains the same DNA—different cell types, such as liver and muscle cells, selectively express the genes for production of characteristic proteins. | Predict the pattern of gene expression expected for different genes based on the cell type. |
| Epigenetics |  | Differentiate between an endocrine disruptor and a mutagen in terms of gene expression in a cell. |
| **Ecology Recommended Learning Objectives** |  |  |
| **Topic** | **LOCS** | **HOCS** |
| Climate change | Explain the biology behind policies to limit carbon emissions in order to minimize impacts of climate change. | Given a list of personal changes, evaluate the contribution of each to meaningfully impact climate change. |
| Climate change |  | Compare features that are important inputs and outputs in a climate change model. |
| Climate change |  | Evaluate policies to limit carbon emissions in order to minimize impacts of climate change. |
| Climate change |  | Given data, identify and test a hypothesis about how a population will respond to climate change. |
| C-cycles | Explain changes in the carbon cycle including the role of human activity since the industrial revolution. | Make predictions about climate change outcomes based on carbon cycle changes. |
| C-cycles | Describe the carbon cycle in terms of fast and slow carbon, and identify carbon sources and reservoirs. | Predict the impact of the following types of events on atmospheric CO<sub>2</sub> levels: 1) extensive tree planting programs, 2) increases in cellular respiration that occur when warming temperatures increase decomposition rates in the arctic. |
| Alternative Energy & Biofuels | Define common energy sources, including biofuels, and explain where they are sourced from. | Investigate the environmental and economic costs of energy sources including renewable energy sources such as biofuels, and non-renewable chemical energy such as oil and gas. |
| Alternative Energy & Biofuels | Describe how biofuels function as energy sources. |  |
| Alternative Energy & Biofuels | Describe how environmental resource extraction, using examples such as fossil fuel extraction, fracking, and logging, impacts environmental health. |  |
| Biodiversity & Ecosystem Health | Describe human activities that impact biodiversity. | Analyze evidence that there has been a change in global biodiversity over the last 100 years. |
| Biodiversity & Ecosystem Health | Identify the aspects of biodiversity that can be used to determine an ecosystem's value. | Evaluate trade-offs between different stakeholders for an ecosystem, including environmental health, cultural use, and economic value. |
| Biodiversity & Ecosystem Health | Graphically, verbally, or quantitatively describe a population over time. |  |
| Biodiversity & Ecosystem Health | Identify the human health consequences of environmental degradation such as dioxin, water borne diseases, and poor air quality. | Construct an argument that relates a human health issue with the environment (e.g., water-borne disease, food poisoning, air quality, dioxin, UV radiation). |
| Bioremediation & Soil | Explain the role that soil plays in terrestrial ecosystems. |  |
| Bioremediation & Soil | Explain why high levels of organic matter are important in soil. |  |
| Bioremediation & Soil | Identify environmental consequences of agricultural practices such as tilling and fertilizer use. |  |
| Bioremediation & Soil | Explain the biological process of composting. |  |
| Bioremediation & Soil | Describe the benefits of composting as a carbon sink, for agriculture, and for reducing waste. |  |
| Bioremediation & Soil | Explain the role that soil plays in terrestrial ecosystems. |  |
| Sustainable Crops & GMOs | Describe why someone would be concerned about or support genetic modifications in organisms like crops. | Discuss the costs and benefits associated with genetically modified organisms, including human gene therapy. |
| Sustainable Crops & GMOs |  | Distinguish between selective breeding and genetic modifications. |
| Sustainable Crops & GMOs |  | Find and evaluate research studies that serve as the basis for understanding the safety of genetic manipulations in crops, and use these scientific resources to support an a |
| Aquatic Biomes |  | Identify trends in data (e.g., population size and density) on species in a habitat. |
| Aquatic Biomes |  | Analyze the role of organisms as producers, consumers, and decomposers in marine ecosystems. |
| Aquatic Biomes |  | Develop and use a model of the movement of energy in marine ecosystems to evaluate the impact of an environmental perturbation. |
| Aquatic Biomes |  | Justify how water temperature, depth, stratum, location, and/or time of year affect the abundance of organisms at a location. |
| Aquatic Biomes |  | Predict and explain your reasoning about where you would find a specific species in a particular habitat, if given information on the natural history of a species. |
| Aquatic Biomes |  | Compare different perspectives of stakeholders involved in regulating fisheries. |
| Eutrophication | Define “dead zones” and eutrophication. | Predict how increasing nitrogen impacts microbes in an ecosystem. |
| Eutrophication | Identify human behaviors that lead to excess nitrogen in the environment. | Critique approaches to human waste in terms of environmental sustainability. |
| Coral Bleaching | Explain the benefits of the symbiotic relationship between corals and zooxanthellae. | Graph and interpret data to estimate potential harm to an ecosystem. |
| Coral Bleaching | Characterize the locations of coral reef ecosystems around the world. | Analyze a graph to predict the occurrence, timing and severity of coral bleaching. |
| Coral Bleaching | Explain the consequences of rising sea surface temperature on coral reefs. | Calculate accumulated heat stress from sea surface temperature data. |
| Ocean Acidification |  | Predict the likely effects of changes of increased CO<sub>2</sub> on ocean pH |
| Ocean Acidification |  | Given a research question about ocean acidification, gather data to answer it. |
| Plastic Pollution | Describe the environmental challenges of plastic pollution. | Interpret graphs and maps of plastic pollution. |
| Plastic Pollution | Characterize how plastic and other materials should be appropriately recycled. | Evaluate proposed models of plastic pollution reduction. |
| Microbiomes | Explain the role of microbiomes in human health. | Differentiate between mutualism, commensalism, and parasitism and recognize examples of these in human-bacterial symbiotic relationships. |
| Microbiomes | Explain the difference between species diversity and richness and recognize examples of each. |  |
| **Evolution Recommended Learning Objectives** |  |  |
| **Topic** | **LOCS** | **HOCS** |
| Natural Selection | List the critical steps for evolution to occur by natural selection. | Distinguish between the idea that selection acts on individuals but only populations evolve. |
| Antibiotic Resistance | Explain how DNA sequences can be used to trace the evolution and spread of antibiotic resistance. |  |
| Antibiotic Resistance | Provide examples of antibiotic-resistant bacteria and explain why antibiotic resistance is a problem. |  |
| **Genetics Recommended Learning Objectives** |  |  |
| **Topic** | **LOCS** | **HOCS** |
| Inheritance | Define the following terms: CHROMOSOME--and distinguish replicated and unreplicated--CHROMATID, PLOIDY, HAPLOID NUMBER, HOMOLOGOUS CHROMOSOMES | Compare the genetic information held on two homologous chromosomes, two nonhomologous chromosomes and two sister chromatids. |
| Inheritance | Explain how the combination of alleles determines phenotype for a simple monogenic trait with a dominant/recessive pattern of inheritance | Calculate the probability of a particular gamete being produced from an individual, assuming independent segregation. |
| Inheritance | Label which elements in a Punnett square represent the genotypes of egg, sperm, and offspring. Explain how you can determine the frequency of each egg and sperm | Calculate the probability of a particular genotype, given independent segregation and random union of gametes between two individuals with known genotypes. |
| Inheritance |  | Given any pair of parental genotypes and information on the alleles present, use a Punnett square to complete a genetic cross. Identify the genotypes and phenotypes of offspring and calculate their predicted frequencies. Note that the genes involved may be autosomal, X-linked, linked, or unlinked and that the alleles involved may be dominant, recessive, or co-dominant. |
| Inheritance |  | Given information about two parents, use Punnett squares to determine the probability that a child will inherit (1) an autosomal dominant, (2) an autosomal recessive, or (3) a sex-linked recessive allele and/or show symptoms of the disease caused by this allele if given information about the parents |
| Inheritance |  | Given a completed Punnett square and information on offspring phenotypes, determine whether the alleles involved are 1) dominant, recessive, or codominant, 2) autosomal or X-linked, and 3) linked or unlinked. |
| Inheritance |  | Using pedigrees, distinguish between dominant, recessive, autosomal, X-linked monogenic traits |
| Genetic Variations | Describe different types of genetic variations (mutations) in the genome, including their location in coding and noncoding DNA. | If given information about the type and genome location of an inherited genetic variation, predict whether it will alter the structure or amount of a protein inside cells, and whether the genetic variation is likely to change a trait. |
| Genetic Variations | Explain the difference between chromosomes, genes, DNA, RNA, and proteins | Rank the following mutations in terms of greatest to least impact on the structure and function of genes and gene products: missense (change amino acids), nonsense (change to "stop"), frameshift (change reading frame), and silent (no change in product). Explain your reasoning. |
| Genetic Variations | Evaluate how genes and the environment can interact to produce a phenotype. | Predict how a given mutation in DNA may or may not affect that amino acid sequence of the |
|  |  | protein produced (through transcription and translation) |
| Genetic Variations | Explain how continuous traits involve many different genes, each with alleles that can contribute varying amounts to the phenotype. |  |
| Genetic Testing | Explain the purpose of a genetic test. |  |
| Genetic Testing | Discuss the ethical issues involved in obtaining and using data on DNA sequences and chromosome structure in human parents and fetuses. | Research real-life genetic testing scenarios and consider the advantages and disadvantages associated with common genetic tests for different traits. |
| Genetic Testing | Identify genetic and environmental factors linked to the development of a human trait. | Research genetic and environmental factors linked to the development of a human trait using materials on direct-to-consumer genetic testing sites. |
| Genetic Testing | Describe a study design that examined a link with a human trait and a gene. |  |
| Genetic Testing | Describe the product from a gene linked to a human trait and explain why some alleles are associated with different reactions. |  |
| Non-Biological Basis of Race | Provide examples of the social history of race. | Use evidence to defend the statement that race cannot be defined using biological traits. |
| Non-Biological Basis of Race | Compare the biological definition of genomic ancestry and the social definition of race. | Analyze examples where race is useful and important in biomedical research—and how it is not useful. |
| Non-Biological Basis of Race | Use evidence to explain why light or dark skin color is beneficial or harmful in a particular environment, especially before vitamin D supplements. |  |
| Non-Biological Basis of Race | Use what you know about UV radiation and DNA damage, to discuss the role of melanin, mutations and skin cancer. |  |
| Non-Biological Basis of Race | Explain the causes and consequences of a racial health disparity. | Critique the use of physical traits to define race. |

Faculty rated the following Vision & Change competencies as the three most critical for non-majors to learn: apply evidence-based reasoning and biological knowledge in daily life (e.g., consuming popular media, deciding how to vote); use a variety of modes to communicate science (e.g. oral, written, visual); and analyze data, summarize resulting patterns, and draw appropriate conclusions.

### Delphi Round 2 Survey

In Delphi Survey Round 2 we asked the participating faculty evaluators to evaluate whether each of the 270 candidate LOs would be considered critical in an introductory biology course for non-majors. Most of the lower order LOs were evaluated as critical more than half of the time. However, whether higher order LOs were evaluated as critical was more evenly distributed, indicating that faculty shared lower agreement about the importance of higher order Bloom’s LOs than for lower order LOs (Figure 3). Yet, more than 30 higher order LOs were rated as critical by faculty. Of the LOs evaluated in the Delphi Round 2 survey, 59% included at least one Vision & Change competency, as compared to only 17.7% of instructors’ LOs and 7% of the textbook LOs from a prior analysis of learning objectives obtained from a national survey (Heil, et al., 2023).

**Figure 3.**
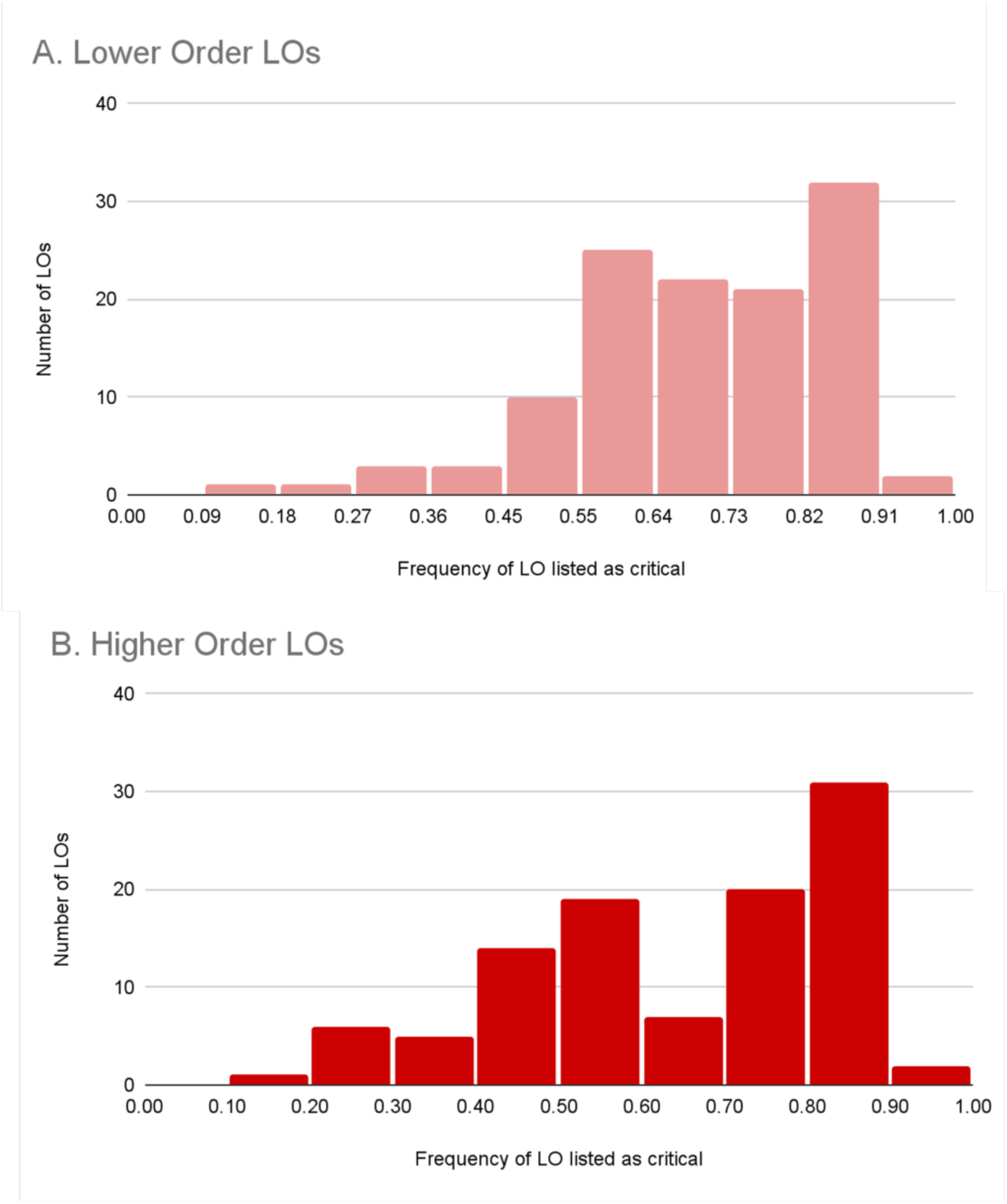
Evaluation of critical level for (a) lower order and (b) higher order LOs.

**Figure 4.**
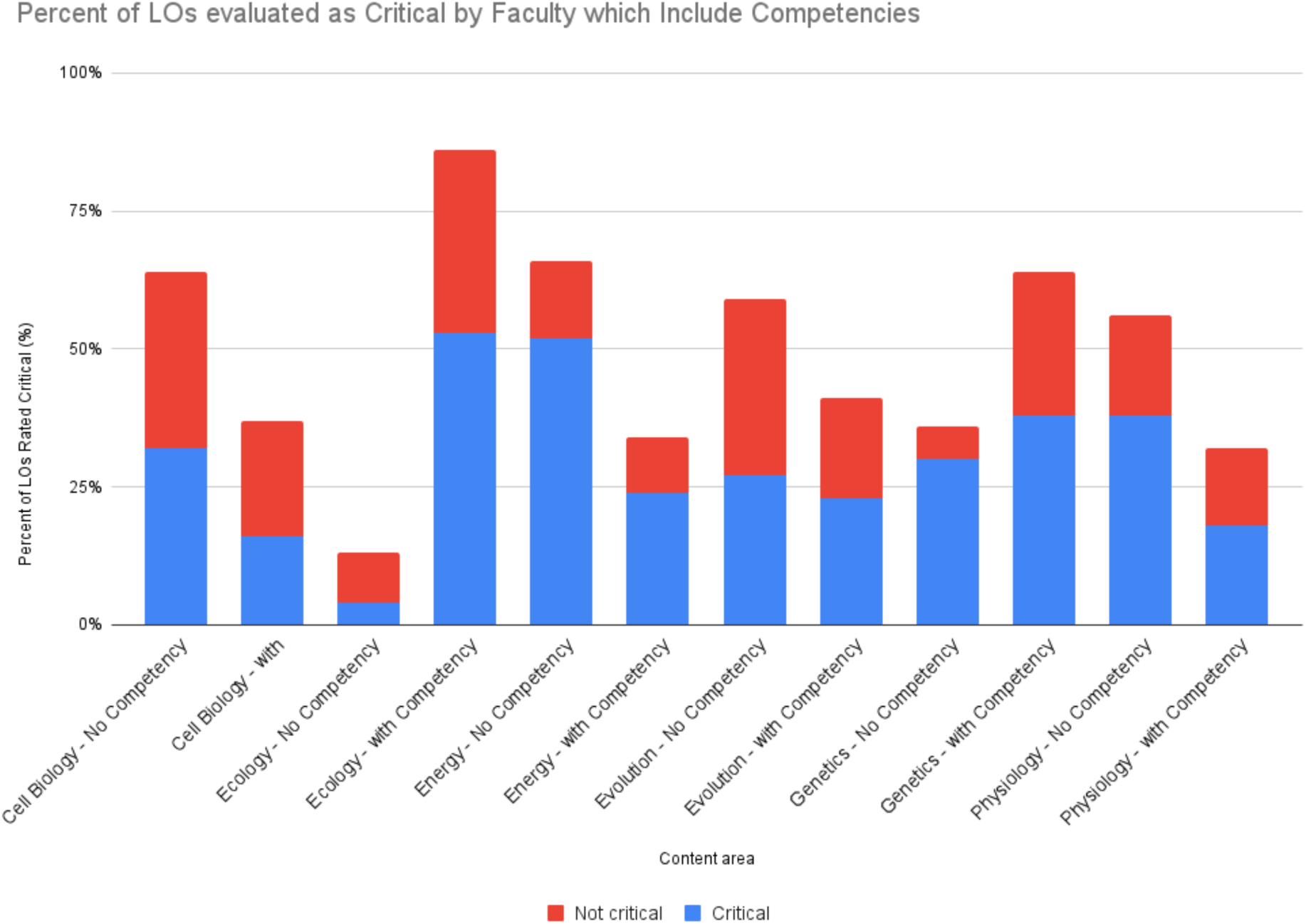
Percent of LOCS and HOCS LOs that were considered critical for each content area.

**Figure 5.**
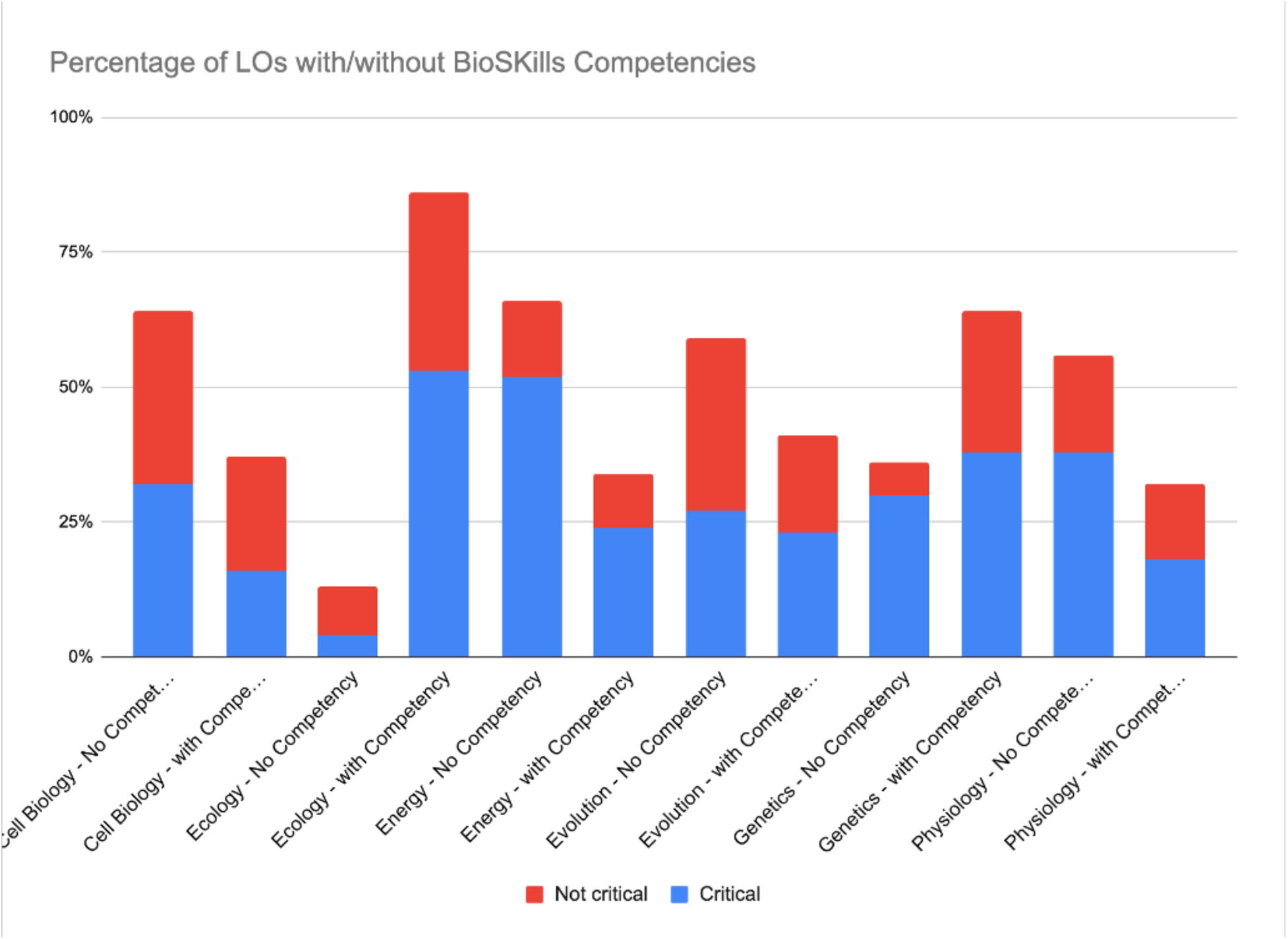
Percent of LOs evaluated as non-critical by faculty which include competencies, categorized by content area. Some LOs include more than one competency. Competency is indicated in the graph, which includes all LOs that assessed either none or at least one BioSkills competency.

### Delphi Round 3 Survey

Once a master dataset of responses to the Delphi Round 2 survey was assembled, we took all the data and created collaborative google sheets for each content area to share with our Delphi panelists in the third round of evaluation. This sheet contained all LOs with higher and lower order indicated as well as the ratio of evaluators who had rated each LO as critical, not critical, and suggested changes. We included statements that indicated which of the LOs have >70% agreement as critical and explained that there was no need to comment on these LOs. We separately binned the LOs that reviewers provided lower agreement for whether they were critical or not. For each LO, we asked the reviewers to: (1) indicate if they thought the LO should be included in our final set of official LOs for non-major’s biology and (2) explain their reasoning.

We examined all comments where participants argued that the LO was critical. The most frequent comments provided by delphi round 3 participants related to: (1) the LO was evaluated as not relevant to student learning about the socio-scientific issue (27%); (2) the LO was evaluated as relevant to student learning about this issue (26%); and (3) the evaluator judged the LO as too detailed (11%) or simply recommended deleting the LO (11%).

### Recommending a set of LOs for non-majors biology

This project’s goal was to develop a set of lesson level LOs for an issues-focused introductory biology course for non-majors as consistent with programmatic goals articulated in the *Vision and Change* report and that have been reviewed by a focused group of expert instructors. Of the 270 original LOs, 60.7% were deemed critical. Expert instructors also recommended 22 additional new LOs. However, these LOs have not undergone this evaluation process. (See Links to Supporting Data).

Since most non-majors courses are confined to one semester, we recommend that faculty select 1-4 issues, and consider using all of the LOs that received high endorsement based on percent-essential ratings. Following these guidelines, we recommend a total of 164 candidate LOs as the core LOs for introductory biology for non-majors courses (Table 3)

## Discussion

This project’s goal was to develop a set of lesson level learning objectives (LOs) for a non-majors introductory biology course that were consistent with programmatic goals articulated in the *Vision and Change* report and endorsed by expert faculty. The project began with 270 novel LOs developed for 14 units, each based on one unique socio-scientific issue, across 6 biological topics. Through the evaluation process, 60.7% were deemed critical, and experts also recommended 22 additional new LOs.

Learning objectives offer a unified framework for faculty and students to make instructional goals clear and can function to document and improve student learning (Orr et al., 2022). Learning objectives can be used to develop formative and summative assessments (Mager & Peatt, 1997). Learning objectives are a critical component of backward course design, necessary for aligning instructional practice with assessment (Fink, 2003). Given recent work about instructional practices to promote science literacy for non-majors, we developed LOs focused around relevant socio-scientific issues (Gormally and Heil, 2022).

Interestingly, expert evaluators rated learning objectives that addressed lower-order cognitive skills (LOCS) as well as higher-order skills (HOCS) as critical. Much research has documented that introductory courses for life sciences majors tend to prioritize LOCS, emphasizing content coverage and memorization rather than HOCS, despite policy recommendations (Derting et al., 2016; Momsen et al., 2010). Indeed, Hennessey & Freeman (2024) report that instructors in majors classes were more likely to rate a LO as essential if it addressed a LOCS. Likewise, recent work by Heil et al. (2024) report an emphasis on LOCS in non-majors syllabi. Our findings are a positive shift that aligns with education policy recommendations.

Education policy continues to call for making science learning useful for students. From research, we know that prioritizing socio-scientific issues in science learning is one critical approach to making science learning useful (Gormally & Heil, 2022). Recent analysis of syllabi from non-majors biology courses indicated that nearly half (48%) focus solely on science content, giving little attention to socio-scientific issues (Heil et al., 2024). Given this background, our LO development prioritized contextualizing science learning across fourteen unique socio-scientific issues. These issues include antibiotic resistance; the role of biology and genetics in our understanding of race; viruses and pandemics; cancer; environmental issues including biodiversity loss and ocean acidification; among other socio-scientific issues. Consequently, these LOs meet the call from Vision & Change to support students “to participate as citizens and thrive in the modern world.”

This project’s goal was to develop a set of lesson level LOs for two semesters of introductory biology for non-majors, consistent with the programmatic goals as articulated in Vision & Change. These LOs have been evaluated by a diverse group of faculty experts. In total, 60.7% of LOs were endorsed as essential by faculty experts. This means a total of 164 finalized LOs out of 270 original LOs were endorsed as LOs to use in introductory biology for non-majors courses.

Given the diverse nature of non-majors biology courses, we recommend instructors select units based on the needs of their particular university, program, course, and student interest. Following Heil et al.’s (2023) work, we would suggest estimating 36 class sessions per semester course, not including exams and vacation days.

Hennessey & Freeman (2024) recommend using three LOs per class session, which suggests a total of 108 LOs may be reasonable for a one semester course, approximately 64.7% of the total LOs available. Further, Hennessey & Freeman (2024) propose that instructors utilize ∼75% of endorsed LOs in addition to ∼25% of their own LOs that reflect their program and student needs.

### Study Limitations

Our study limitations included factors that were influenced by the COVID-19 pandemic as well as subsequent faculty burnout. Recruiting and retaining faculty participants was an ongoing challenge throughout the project. This impacted our sample size, the number of rounds of feedback we conducted, as we saw faculty fatigue from participation in research. In a 2020 Chronicle of Higher Education study of 1,122 faculty, more than half reported considering retiring or changing careers from academia (Education, 2020). In a study of 530 full- and part-time faculty entitled, “Burnt Out and Overburdened: The Faculty Experience, 2022,” 1 in 2 faculty members reported burnout (O’Donnell, 2023). Among the 530 faculty, 55% reported not having enough time to teach effectively and of those reporting burnout, 72% reported considering leaving higher education. Given this context, it is perhaps unsurprising that recruiting and retaining faculty participants was one challenge we faced throughout this project. It is likely that faculty burnout may be higher amongst faculty who teach at less-resourced colleges and universities. Consequently, this may have impacted the representation of faculty from two-year institutions, masters colleges and universities and baccalaureate institutions. Finally, unrelated to the pandemic and faculty burnout, we faced the additional challenge that as a field, we continue to lack a consensus about what non-majors should learn and we prioritize majors’ learning.

### Future work & conclusions

The LOs including rankings and presence of competencies have been shared with Howard Hughes Medical Institute (HHMI) and will be included in the HHMI assessment builder. Additionally, these LOs can be used as models by other faculty and instructional designers developing LOs focused on competencies. Future work may focus on the development of novel curricula including CUREs and service-learning opportunities for students to practice the LOs in high impact, authentic learning experiences. Future research questions may focus on assessing student learning using the LOs, as well as exploring student interest and engagement in learning about specific content areas and socio-scientific issues.

## Acknowledgements

We thank our faculty research participants for their time. Without their contributions, this work would not be possible. We appreciate the critical feedback we received from Austin Heil, Scott Freeman, Kelly Hennessey, Melissa Csikari, and Alexa Clemmons.

This work was supported by funding from the National Science Foundation IUSE Award #2012362.

