## Supplemental Materials for "Development of Learning Objectives for Non-Major Introductory Biology Using a Delphi Method"

**Table 1.** Description of content areas related to specific socioscientific issues

| Unit & Socioscientific issue | Description |
|---|---|
| Biochemistry | |
| Macromolecules & diet, poisons & metabolic pathways | Macromolecules, nutrition and food labels, metabolic pathways and enzymes |
| Cell Biology | |
| Cancer & the Cell Cycle, Gene Expression, Epigenetics, and Food Deserts | DNA and RNA; Societal implications of food deserts, chronic stress and multigenerational impacts; diet and nutrition; regulating gene expression; estrogen, endocrine disruptors, and the environment; cancer and the epigenome |
| Ecology | |
| Climate change, C-cycles, biofuels | Biodiversity, indigenous culture, biogeochemical cycles, and biofuels |
| Biodiversity loss, Ecosystem Health, Bioremediation & Soil, GMOs & sustainable food | Ecosystems and habitat loss; growing food, soil remediation, phytoremediation and pollution, GMOs & sustainable food, and food insecurity |
| Overfishing, Eutrophication, Coral Bleaching, Ocean acidification, plastic pollution | Food chains; biogeochemical cycles; habitats; biodiversity; overfishing, eutrophication, coral bleaching, ocean acidification, plastic pollution |
| Evolution | |
| Antibiotic Resistance, Microbiomes, Phylogenetics | Drug-resistant infections; natural selection; microbiomes, and foodborne illnesses |
| Genetics | |
| Inheritance and Genetic Testing | Genetic variation and mutations; genetic testing techniques (PCR, gel electrophoresis, gene sequencing); patterns of Inheritance, pedigree analysis; complex genetic traits |
| How race is and is not biological | Social constructs of race; genomic ancestry; human migration out of Africa; melanin and UV radiation; racial health disparities |

| Animal Physiology | |
|---|---|
| Immune System, Pandemics, HIV, & Vaccines | Vaccination and the anti-vaxxer movement; personal and herd immunity; allergies; and evolution of viruses |
| Nervous system, stress, anxiety and depression | How nervous systems work; stress responses; pain; and how antidepressants work |
| Sex, reproduction, & gender | Inclusive sex, gender, sexual orientation, reproduction; human fertility; how hormones for transitioning and birth control work |
| Pain, human sensory experiences, disability | Sensory systems and biological, medical, and societal perspectives about dis/ability |

**Table 2.** Coding of faculty evaluator comments about LOs and example comments.

| Code | Definition | Example Quote | Percent of comments (%) |
|---|---|---|---|
| not relevant | Reviewer states that this LO is not relevant to real-world issues that non-majors need to learn | No, I can't think of why students would need to be able to do this in their lives. I can't think of how this information would be useful. | 27 |
| relevant/critical | Reviewer states the LO is relevant to students' real-world learning | Yes. This is quite germane for any and all student populations. In my many years of teaching this topic, I have been struck by how much misinformation about this topic is in the heads of students (for lots of reasons, not blaming or shaming here). They love learning about this. | 26 |
| delete LO | Reviewer recommends deleting the LO | I will just generally comment for all of these that I agree they can be omitted. Given the breadth of potential LOs on these topics, these specific ones are not critical to include. | 11 |
| Too detailed | Reviewer states that the LO is too detailed for non-majors | This feels too specific. There are so many cancer treatments out there, I don't want students to focus on one type. | 11 |

| | | | |
|---|---|---|---|
| how to use LO | Reviewer provides feedback about how to use the LO | Yes, if it's used as more of an application of the other objectives; once they understand cycles, etc., then identify how variations can cause infertility. | 10 |
| too difficult for non-majors | Reviewer states that this LO is too difficult or confusing for non-majors to learn | For a non-majors class this might be too difficult, especially if you have not talked about analyzing sequences | 5 |
| too vague | Reviewer states LO is too vague | I am not sure what this means. I need more information. Microbiomes where? On Earth, in our bodies? I love having students analyze data, but what is the question being addressed with the data? | 3 |
| keep LO | Reviewer recommends keeping LO | I would include both of these | 1 |
| skill is needed | Reviewer states that students need to develop a skill for this LO | For all of these, I like the higher-order skill focus but think that they are too specific to be considered essential. I would be on board with a more general "draw conclusions from data about cancer risk and/or cancerous cellular activity" and allow instructors to take that in directions that fit the analysis tools and specific topics they've emphasized in class. | 1 |
| not critical here | Reviewer states LO is critical but not for this unit | I think it's important for students to understand synthetic genetic manipulation and more traditional breeding programs (and that they're both GMOs), but that is likely captured elsewhere. | 1 |
| redundant | Reviewer states LO is redundant | I think this one is captured in the first LO, but I think both of the above are great LOs for understanding societal challenges and how their bodies process pain. | 1 |

**Table 3.** Ranked Importance of Socioscientific Issues for Non-majors Biology

| Issue | Number of times mentioned in the top 5 | Probability of being in the top 5 per each faculty | Round 2 Delphi LOs | |
|---|---|---|---|---|
| | | | LOCS / HOCS | % critical |
| Climate change, C-cycles, biofuels | 16 | 80.0% | 42% LOCS | 67% |
| | | | 58% HOCS | 56% |
| Apocalyptic pandemics - Immunity | 15 | 75.0% | 60% LOCS | 47% |
| | | | 40% HOCS | 70% |
| How race is and is not biological | 13 | 65.0% | 38% LOCS | 80% |
| | | | 62% HOCS | 50% |
| Antibiotic Resistance and Microbiomes | 13 | 65.0% | 73% LOCS | 63% |
| | | | 27% HOCS | 17% |
| Sustainable food choices & GMOs | 8 | 40.0% | 44% LOCS | 71% |
| | | | 56% HOCS | 78% |
| Gene expression, epigenetics, cancer, and diet | 8 | 40.0% | 63% LOCS | 33% |
| | | | 37% HOCS | 57% |
| How genetic testing works | 5 | 25.0% | 47% LOCS | 78% |
| | | | 53% HOCS | 60% |
| Ocean ecosystems, plastic pollution, and climate change | 5 | 25.0% | 20% LOCS | 50% |
| | | | 80% HOCS | 88% |
| Sex verification tests and what biology tells us sex and gender | 5 | 25.0% | 50% LOCS | 100% |
| | | | 50% HOCS | 40% |
| Stress, anxiety, depression and the nervous system | 5 | 25.0% | 78% LOCS | 64% |
| | | | 22% HOCS | 50% |
| Understanding human sensory | 2 | 10.0% | 73% LOCS | 91% |

| | | | | |
|---|---|---|---|---|
| experiences, pain, & disability | | | 27% HOCS | 50% |
| Genetic variation of alcohol metabolism, poisons, & metabolic pathways | | | 64% LOCS | 71% |
| | 0 | 0.0% | 36% HOCS | 75% |
| TOTALS | 95 | from 20 faculty | | |

**Table 4.** Vision & Change Competencies for Non-majors Biology. Faculty were asked to rank the importance of the competencies.

| **Process of Science** | **Times Ranked in top 5** | **Round 2 Delphi LOs which include the competency** |
|---|---|---|
| Analyze data, summarize resulting patterns, and draw appropriate conclusions. | 14 | 27 (10%) |
| Design controlled experiments including plans for analyzing the data. | 1 | 3 (1%) |
| Formulate testable hypotheses and state their predictions | 3 | 2 (1%) |
| **Quantitative Reasoning** | | |
| Create and interpret informative graphs and other data visualizations. | 9 | 2 (0.7%) |
| Interpret the biological meaning of quantitative results. | 8 | 15 (6%) |
| Perform basic calculations (e.g., percentages, frequencies, rates, means). | 1 | 8 (3%) |
| Select and apply appropriate equations (e.g. Hardy- Weinberg, Nernst Gibbs free energy) to solve problems. | 0 | 0 |
| Use rough estimates informed by biological knowledge to check quantitative work | 0 | 0 |
| **Modeling** | | |
| Build and revise conceptual models to propose how a biological system or process works | 3 | 6 (2%) |
| Identify important components of a system and describe how they influence each other (e.g. positively or negatively) | 4 | 26 (10%) |
| Summarize relationships and trends that can be inferred from a given model or simulation | 2 | 18 (7%) |
| Use models and simulations to make predictions and refine hypotheses, note similarity to process of science statement in | 3 | 11 (4%) |

| | | |
|---|---|---|
| hypotheses and predictions | | |
| **Interdisciplinary Nature of Science** | | |
| Build models or explanations of simple biological processes that include concepts from other STEM disciplines or multiple fields of biology. | 4 | 1 (0.3%) |
| Describe examples of real-world problems that are too complex to be solved by applying biological approaches alone. | 7 | 15 (6%) |
| **Communication and Collaboration** | | |
| Use a variety of modes to communicate science (e.g. oral, written, visual). | 15 | 14 (5%) |
| **Science and Society** | | |
| Identify and describe the broader societal impacts of biological research on different stakeholders. | 12 | 41 (15%) |
| Apply evidence-based reasoning and biological knowledge in daily life (e.g., consuming popular media, deciding how to vote). | 15 | 11 (4%) |
| **Totals** | N = 20 100 responses | N = 270 LOs |

#### Round 1 Survey - What is important for non-science majors to learn?

Your syllabus has identified your non-majors biology course as exemplary. In this survey, we are interested in what socioscientific issues (i.e., controversial, socially relevant, real-world problems that are informed by science) and competencies (i.e., skills) you find most important for your non-science-major students. You will be provided space to explain your choices and add issues that you think we might have missed.

##### Part 1: Socioscientific issues

1. Among the list below, please select the TOP FIVE socioscientific issues (controversial, socially relevant, real-world problems that are informed by science) you find most important to non-majors biology courses. Issues can be found in manuscript
   a. Apocalyptic pandemics: vaccination and the anti-vaxxer movement; personal and herd immunity; and evolution of viruses
   b. How race is and is not biological: social constructs of race; genomic ancestry; human migration out of Africa; melanin and UV radiation; racial health disparities
   c. Stress, anxiety, and depression: how nervous systems work; stress responses; and how antidepressants work

d. Sustainable food choices & GMOs: macromolecules, nutrition and food labels, ecosystems and habitat loss; natural and artificial selection and evolution; food insecurity
e. Genetics of alcoholism: genetic variation of alcohol metabolism; metabolic pathways and enzymes; transcription and translation
f. Climate change, environmental resources and conflicts: biodiversity, indigenous culture, biogeochemical cycles, and biofuels
g. How genetic testing works: DNA and RNA; genetic variation and mutations; genetic testing techniques (PCR, gel electrophoresis, gene sequencing); patterns of Inheritance, pedigree analysis; complex genetic traits
h. How ocean ecosystems are impacted by plastic pollution and climate change: food chains; biogeochemical cycles; habitats; biodiversity
i. Sex verification tests and what biology tells us sex and gender: inclusive sex, gender, sexual orientation, reproduction; how hormones for transitioning and birth control work
j. Understanding human sensory experiences through a diversity lens: sensory systems and biological, medical, and societal perspectives about dis/ability
k. Antibiotic resistance and the microbiome: drug-resistant infections; natural selection; foodborne illnesses
l. Food waste and composting: growing food, soil remediation, phytoremediation and pollution
m. Gene expression, epigenetics, cancer, and diets: societal implications of food deserts, chronic stress and multigenerational impacts; diet and nutrition; regulating gene expression; estrogen, endocrine disruptors, and the environment; cancer and the epigenome

2. Click and drag each item to rank your top five socioscientific issues, with 1 representing the most important for non-major courses. (Question answers populated with the 5 responses from the last question).
3. Explain how you selected and ranked your top socioscientific issues.
4. Please tell us socioscientific issues that you think are important for non-majors that we are missing. Click here for our complete list.

#### Part 2: Competencies

5. Rank the following competencies in terms of importance for non-science majors to learn.

Process of Science

a) Analyze data summarize resulting patterns and draw appropriate conclusions.
b) Design controlled experiments including plans for analyzing the data.
c) Formulate testable hypotheses and state their predictions.

Quantitative Reasoning

a) Create and interpret informative graphs and other data visualizations.
b) Interpret the biological meaning of quantitative results.
c) Perform basic calculations (e.g. percentages, frequencies, rates, means).
d) Select and apply appropriate equations (e.g. Hardy- Weinberg, Nernst Gibbs free energy) to solve problems.
e) Use rough estimates informed by biological knowledge to check quantitative work.
    a. Modeling

f) Build and revise conceptual models to propose how a biological system or process works.
g) Identify important components of a system and describe how they influence each other (e.g. positively or negatively).
h) Summarize relationships and trends that can be inferred from a given model or simulation.
i) Use models and simulations to make predictions and refine hypotheses, note similarity to process of science statement in hypotheses and predictions.

Interdisciplinary Nature of Science

a) Build models or explanations of simple biological processes that include concepts from other STEM disciplines or multiple fields of biology.
b) Describe examples of real-world problems that are too complex to be solved by applying biological approaches alone.
    a. Communication & Collaboration
c) Use a variety of modes to communicate science (e.g. oral, written, visual).
    a. Science & Society
d) Identify and describe the broader societal impacts of biological research on different stakeholders.
e) Apply evidence-based reasoning and biological knowledge in daily life (e.g., consuming popular media, deciding how to vote).

6. Click and drag each item to rank your top five competencies, with 1 representing the most important for non-major courses.
7. Explain how you selected and ranked your top competency skills. (open-response)
8. Please tell us competencies that you think are important for non-majors that we are missing. Click here for our complete list of competencies. (open-response)

#### Part 3 Brief Survey

Please rate your level of confidence for the following two questions.

9. How confident do you feel in your ability to alter your course to fit a set of agreed-upon LOs? Not confident at all; slightly confident; somewhat confident; fairly confident; very confident
10. How likely is your institution to allow you to make changes to your non-science major course? Very unlikely, unlikely, neutral, likely, very likely

#### Part 4 Demographic Questions (Open-response)

First and Last Name
Gender (or state that you prefer not to respond)
Race/Ethnicity (or state that you prefer not to respond)
Number of years teaching (total)
Number of years teaching non-science majors
Institution Name
Institution Type (select all that apply) PhD-granting, MS granting, 4-year, minority-serving, liberal arts college, community-college/2-year
Position (select all that apply) Adjunct, tenured/tenure track/non tenure-track, department chair/program head

#### Round 2 Survey - Provide Feedback on an Initial List of Non-Majors Biology Learning Objectives

Our process for crafting the Learning Objectives for non-majors included beginning with an analysis of the content and issues displayed in exemplary syllabi like the one you provided from your course. As the next step in the process, we are interested in your feedback on the learning objectives for different issues commonly taught in non-majors Biology. We have developed Learning Objectives for 12 units. It should take 5-10 minutes to read and provide feedback on each unit (containing 12-45 learning objectives each). We would love it if you could provide feedback on 2-3 units.

We ask that you complete the following demographic questions so that we can determine if we are gathering feedback from a representative population. We will not link your responses with any individual identifying information when sharing the results of this survey.

Please skip to the bottom of this section if you are returning to the survey and have already provided us with this information.

1. What is the name of your current institution?
2. What is or was the focus of your graduate training? (Please select all that apply.)
    a. Molecular/Cellular/Developmental Biology
    b. Anatomy and Physiology
    c. Ecology/Evolutionary Biology
    d. Discipline-Based Education Research
    e. Other (Please Specify)
3. In an average academic year when you are teaching, which levels do you teach? (Please select all that apply.)
    a. Non-Majors Lower-Level (100 - 200 Level)
    b. Mixed-Majors Lower-Level (100 - 200 Level)
    c. Majors Lower-Level (100 - 200 Level)
    d. Upper-Level (300 - 400 Level)
    e. Graduate-Level (500+ Level)
4. To what extent do you communicate learning objectives to your students in your introductory biology course? (Please select all that apply.)
    a. Every class session
    b. Weekly
    c. Unit overview
    d. Course overview
    e. Syllabus
    f. Other (Please Briefly Describe)
5. In a typical academic term when you are teaching in an Introductory Biology for Non-Majors series, what is the focus of the course you teach? (Please select all that apply.)
    a. Biochemistry
    b. Cell Biology
    c. Genetics
    d. Evolution
    e. Biodiversity of Life

    f. Plant and Animal Physiology
    g. Ecology
    h. Other (Please Specify)
6. Do you have Biology Education Research experience?
    a. Yes
    b. No
7. To what extent have the Vision and Change Core Concepts changed your Introductory Biology course design since the report was issued by the AAAS in 2011?
    a. A great deal
    b. Some
    c. Very Little
    d. None
8. To what extent have the Vision and Change Core Competencies changed your Introductory Biology course design since the report was issued by the AAAS in 2011?
    a. A great deal
    b. Some
    c. Very Little
    d. None
9. In a typical academic term when you are teaching in an Introductory Biology for Non-Majors series, what are the primary issues that you focus on in the course you teach? (Please select all that apply.)
    a. Cell Biology (Cancer & the Cell Cycle, Gene Expression, Epigenetics, and Food Deserts)
    b. Biochemistry (Macromolecules & Diet)
    c. Ecology (Climate change, C-cycles, biofuels)
    d. Ecology (Biodiversity loss, soil, GMOs, & sustainable food)
    e. Ecology (Overfishing, Eutrophication, Ocean acidification, plastic pollution)
    f. Evolution (Antibiotic Resistance, Microbiomes, Phylogenetics)
    g. Genetics (Inheritance & Genetic Testing)
    h. Genetics (Non-biological basis of race)
    i. Physiology (Human sensory experiences, disability)
    j. Physiology (Immune System, Pandemics, HIV, vaccines)
    k. Physiology (Nervous system and stress, anxiety, and depression)
    l. Physiology (Sex, reproduction, & gender)
    m. Other (please specify)

10. Each section of this survey has between 12-40 learning objectives for you to read and reflect on. Please select 2-3 of the following that you would like to provide feedback on for this survey today (or as many as you would like that you regularly teach.) If you can only complete a few today, you are welcome to come back and re-take the survey at another point to help review more LOs.
    a. Cell Biology (Cancer & the Cell Cycle, Gene Expression, Epigenetics, and Food Deserts)
    b. Biochemistry (Macromolecules & Diet)
    c. Ecology (Climate change, C-cycles, biofuels)
    d. Ecology (Biodiversity loss, soil, GMOs, & sustainable food)

e. Ecology (Overfishing, Eutrophication, Ocean acidification, plastic pollution)
f. Evolution (Antibiotic Resistance, Microbiomes, Phylogenetics)
g. Genetics (Inheritance & Genetic Testing & Non-biological basis of race)
h. Physiology (Human sensory experiences, disability)
i. Physiology (Immune System, Pandemics, HIV, vaccines)
j. Physiology (Nervous system and stress, anxiety, and depression)
k. Physiology (Sex, reproduction, & gender)
l. Other (please specify)

Q2.1 This block of 19 learning objectives (LOs) are part of the Cell Biology unit and contain the issues:
CANCER & THE CELL CYCLE
GENE EXPRESSION, EPIGENETICS, AND FOOD DESERTS

You are evaluating the LOs for these socioscientific issues. Consider if the LOs are critical for students to learn and if these LOs encompass what students need to learn about this issue.

Q2.2 Please indicate whether the following LOs are critical or non critical for teaching this issue to non-majors.

| TOPIC: CANCER & THE CELL CYCLE | CRITICAL | NOT CRITICAL |
|---|---|---|
| 1. Explain why cancer is 1) associated with tumor mutations that regulate the cell cycle, and 2) more common in older than younger people. | o | o |
| 2. Describe the importance of cell cycle checkpoints, and what triggers each phase. | o | o |
| 3. Predict the consequences of events that alter one or more cell cycle checkpoints. | o | o |
| 4. Explain how changes to the epigenome, for example, methylation, can be a potential trigger for cancer. | o | o |
| 5. Describe how chemotherapy drugs might take advantage of methylation to reactivate gene expression. | o | o |
| 6. If given a microarray, be able to explain what color spots you would expect to see for genes that are expressed in normal cells, or overexpressed or underexpressed in cancer cells. | o | o |

Q2.4 Thinking about the socioscientific issue of CANCER & THE CELL CYCLE, is there anything students need to learn about it that is missing in this list of LOs?

o Yes
o No
Q2.5 Please explain what LOs are missing from the socioscientific issue of CANCER & THE CELL CYCLE.

Q2.6 Please indicate whether the following LOs are critical or not critical for teaching this issue to non-majors.

| GENE EXPRESSION, EPIGENETICS, AND FOOD DESERTS | CRITICAL | NOT CRITICAL |
|---|---|---|
| 1. Identify the location of hydrogen bonds and explain their role in stabilizing the double helix. | o | o |
| 2. Define complementary base pairing. | o | o |
| 3. Predict the sequence of a complementary strand of DNA when given one strand. | o | o |
| 4. Compare the structures and components in DNA from RNA. | o | o |
| 5. Describe the flow of information in cells from gene to protein including the roles of mRNA, DNA, polymerase, promoter, gene, amino acids, proteins, and ribosomes in transcription and translation. | o | o |
| 6. Add elements to your central dogma model that represent 1) rRNA, tRNA, and "other RNAs", 2) DNA replication | o | o |
| 7. If given a strand of DNA, be able to predict the RNA transcript that is produced during transcription and the protein produced during translation. | o | o |
| 8. Describe the structure of chromatin, including the differences between the naked double helix, nucleosomes, histones, and chromosomes. | o | o |
| 9. Compare the condensed and de-condensed states of chromatin, including the role of histones, nucleosomes, DNA methylation and histone modification in changing the state of chromatin. | o | o |
| 10. Describe how—although every cell contains the same DNA—different cell types, such as liver and muscle cells, selectively express the genes for production of characteristic proteins. | o | o |

| | | |
|---|---|---|
| 11. Predict the pattern of gene expression expected for different genes based on the cell type. | o | o |
| 12. Define endocrine disruptors. | o | o |
| 13. Differentiate between an endocrine disruptor and a mutagen in terms of gene expression in a cell. | o | o |

Q2.7 Thinking about the socioscientific issue of GENE EXPRESSION, EPIGENETICS, AND FOOD DESERTS, is there anything students need to learn about it that is missing in this list of LOs?
o Yes
o No

Q2.8 Please explain what LOs are missing from the socioscientific issue of GENE EXPRESSION, EPIGENETICS, AND FOOD DESERTS.
**End of Block: Cell Biology**

**Start of Block: Biochemistry (Macromolecules & Diet)**
Q3.1 This block of 21 learning objectives is part of the Biochemistry unit and contain the issues:
MACROMOLECULES & DIET
POISONS & METABOLIC PATHWAYS

Please note that the learning objectives you are evaluating are for this a socioscientific issue. Consider if they are important for students to learn and if these LOs encompass what students need to learn about this issue.
Q3.2 Please indicate whether the following LOs are critical or not critical for teaching this issue to non-majors.

| **TOPIC: MACROMOLECULES & DIET** | **CRITICAL** | **NOT CRITICAL** |
|---|---|---|
| 1. Identify the repeating basic unit of carbohydrates, lipids, proteins, and nucleic acids. | **o** | **o** |
| 2. Identify the major function of macromolecules. | **o** | **o** |
| 3. Identify an example of a monosaccharide, disaccharide, and polysaccharide. | **o** | **o** |
| 4. Identify structures of fats, phospholipids, and steroids. | **o** | **o** |
| 5. Given an image of a novel lipid, distinguish it as an unsaturated or saturated fat, phospholipid, or steroid. | **o** | **o** |
| 6. Describe at least 3 of the many different functions that proteins serve in cells. | **o** | **o** |

| | | |
|---|---|---|
| 7. Explain the connections between the following statements: 1) amino acids vary widely in size and chemical properties, 2) the shapes of proteins are extremely diverse, and 3) proteins serve a wide array of functions in cells. | **o** | **o** |
| 8. Using knowledge of various food molecules, identify ingredients that contribute protein, carbohydrates, and fat. | **o** | **o** |
| 9. Using a food label, calculate the calories contributed by carbs, fats, and proteins. | **o** | **o** |
| 10. Use the nutritional characteristics on a food label to evaluate the health qualities of your meal. | **o** | **o** |

Q3.3 Thinking about the socioscientific issue of MACROMOLECULES & DIET, is there anything students need to learn about it that is missing in this list of LOs?
o Yes
o No
Q3.4 Please explain what LOs are missing from the socioscientific issue of MACROMOLECULES & DIET.

Q3.5 Please indicate whether the following LOs are critical or not critical for teaching this issue to non-majors.

| TOPIC: POISONS & METABOLIC PATHWAYS | CRITICAL | NOT CRITICAL |
|---|---|---|
| 1. Describe characteristics of enzymes that affect their ability to function. | o | o |
| 2. Explain activation energy and what happens when an enzyme catalyzes a reaction. | o | o |
| 3. Identify a substrate, product, and enzyme for a given reaction. | o | o |
| 4. Predict what would happen to the levels of substrates and products if an enzyme was not working. | o | o |
| 5. Describe the different types of inhibition and regulation of enzymatic reactions. | o | o |

| | | |
|---|---|---|
| 6. Provide examples of cells that use many metabolic pathways and cells that use only a subset of metabolic pathways. | o | o |
| 7. Compare how carbohydrates, proteins, and fats are stored or burned for fuel. | o | o |
| 8. Identify the functions of the organs involved in human digestion. | o | o |
| 9. Apply knowledge of converging metabolic pathways to predict how poisons will work. | o | o |
| 10. Explain how ATP is used by the cell as an energy source. | o | o |
| 11. Use a model of variation in metabolic pathways to predict genotypes that are most likely to have specific phenotypes. | o | o |

Q3.6 Thinking about the socioscientific issue of POISONS & METABOLIC PATHWAYS, is there anything students need to learn about it that is missing in this list of LOs?
o Yes
o No
Q3.7 Please explain what LOs are missing from the socioscientific issue of POISONS & METABOLIC PATHWAYS.
**End of Block: Biochemistry (Macromolecules & Diet)**

**Start of Block: Ecology (Climate change, C-cycles, biofuels)**
Q4.1 This block of 19 learning objectives is part of the Ecology unit and contain the issues:
CLIMATE CHANGE
CARBON CYCLES
BIOFUELS
Q4.2 Please indicate whether the following LOs are critical or not critical for teaching this issue to non-majors.

| TOPIC: CLIMATE CHANGE | CRITICAL | NOT CRITICAL |
|---|---|---|
| 1. Interpret the results of an online calculation of a carbon footprint, using an online calculator. | o | o |
| 2. Identify personal changes to meaningfully impact climate change. | o | o |

| | | |
|---|---|---|
| 3. Given a list of personal changes, evaluate the contribution of each to meaningfully impact climate change. | o | o |
| 4. Compare features that are important inputs and outputs in a climate change model. | o | o |
| 5. Explain the biology behind policies to limit carbon emissions in order to minimize impacts of climate change. | o | o |
| 6. Evaluate policies to limit carbon emissions in order to minimize impacts of climate change. | o | o |
| 7. Compare costs and benefits of global efforts to control climate change. | o | o |
| 8. Given appropriate data, calculate the relative carbon footprint of various food items. | o | o |
| 9. Given data, identify and test a hypothesis about how a population will respond to climate change. | o | o |

Q4.3 Thinking about the socioscientific issue of CLIMATE CHANGE, is there anything students need to learn about it that is missing in this list of LOs?
o Yes
o No

Q4.4 Please explain what LOs are missing from the socioscientific issue of CLIMATE CHANGE.
Q4.5 Please indicate whether the following LOs are critical or not critical for teaching this issue to non-majors.

| TOPIC: CARBON CYCLES | CRITICAL | NOT CRITICAL |
|---|---|---|
| 1. Given the summary reactions for photosynthesis and cellular respiration, compare and contrast their reactants and products. | o | o |
| 2. Predict the impact of the following types of events on atmospheric CO2 levels: 1) extensive tree planting programs, 2) increases in cellular respiration that occur when warming temperatures increase decomposition rates in the arctic. | o | o |

| | | |
|---|---|---|
| 3. Given environmental conditions, predict whether photosynthesis and/or cellular respiration is occurring and its role in rising atmospheric CO2. | o | o |
| 4. Describe the carbon cycle in terms of fast and slow carbon, and identify carbon sources and reservoirs. | o | o |
| 5. Describe what characteristics of an ecosystem make it a carbon reservoir. | o | o |
| 6. Explain changes in the carbon cycle including the role of human activity since the industrial revolution. | o | o |
| 7. Make predictions about climate change outcomes based on carbon cycle change | o | o |

Q4.6 Thinking about the socioscientific issue of CARBON CYCLES, is there anything students need to learn about it that is missing in this list of LOs?
o Yes
o No

Q4.7 Please explain what LOs are missing from the socioscientific issue of CARBON CYCLES.
Q4.8 Please indicate whether the following LOs are critical or not critical for teaching this issue to non-majors.

| TOPIC: BIOFUELS | CRITICAL | NOT CRITICAL |
|---|---|---|
| 1. Define common energy sources, including biofuels, and explain where they are sourced from. | o | o |
| 2. Predict the impact of the following types of events on atmospheric CO2 levels: 1) extensive tree planting programs, 2) increases in cellular respiration that occur when warming temperatures increase decomposition rates in the arctic. | o | o |
| 3. Describe how biofuels function as energy sources. | o | o |
| 4. Investigate the environmental and economic costs of energy sources including renewable energy sources such as biofuels, and non-renewable chemical energy such as oil and gas. | o | o |

| | | |
|---|---|---|
| 5. Describe how environmental resource extraction, using examples such as fossil fuel extraction, fracking, and logging, impacts environmental health. | o | o |

Q4.9 Thinking about the socioscientific issue of BIOFUELS, is there anything students need to learn about it that is missing in this list of LOs?
o Yes
o No

Q4.10 Please explain what LOs are missing from the socioscientific issue of BIOFUELS.
**End of Block: Ecology (Climate change, C-cycles, biofuels)**

**Start of Block: Ecology (Biodiversity loss, soil, GMOs, sustainable food)**
Q5.1 This block of 38 learning objectives is part of the Ecology unit and contain the issues:
BIODIVERSITY LOSS
ECOSYSTEM HEALTH
BIOREMEDIATION & SOIL
SUSTAINABLE FOOD & GMOs

Please note that the learning objectives you are evaluating are for this a socioscientific issue. Consider if they are important for students to learn and if these LOs encompass what students need to learn about this issue**.**
Q5.2 Please indicate whether the following LOs are critical or not critical for teaching this issue to non-majors.

| TOPIC: BIODIVERSITY LOSS | CRITICAL | NOT CRITICAL |
|---|---|---|
| 1. Analyze evidence that there has been a change in global biodiversity over the last 100 years. | o | o |
| 2. Describe human activities that impact biodiversity. | o | o |
| 3. Identify the aspects of biodiversity that can be used to determine an ecosystem's value. | o | o |
| 4. Create an environmental impact statement or restoration ecology plan for a specific ecosystem. | o | o |
| 5. Predict and explain the response to disturbance at the level of individual, population, and community. | o | o |

| | | |
|---|---|---|
| 6. Compare a species with a broad range and one with a narrow range. | o | o |
| 7. Graphically, verbally, or quantitatively describe a population over time. | o | o |
| 8. Create a model explaining how populations become better adapted to their environment. | o | o |
| 9. Describe the natural history and important characteristics, for example, reproductive timing, habitat and range requirements of a species. | o | o |
| 10. Use multiple representations such as graphs, equations, diagrams to explain how populations are interacting with abiotic factors. | o | o |
| 11. Describe a census of a population in terms of distribution, size, and density. | o | o |

Q5.3 Thinking about the socioscientific issue of BIODIVERSITY LOSS, is there anything students need to learn about it that is missing in this list of LOs?
o Yes
o No
Q5.4 Please explain what LOs are missing from the socioscientific issue of BIODIVERSITY LOSS.

Q5.5 Please indicate whether the following LOs are critical or not critical for teaching this issue to non-majors.

| TOPIC: ECOSYSTEM HEALTH | CRITICAL | NOT CRITICAL |
|---|---|---|
| 1. If given an ecosystem, describe its most important characteristics. | o | o |
| 2. Evaluate trade-offs between different stakeholders for an ecosystem, including environmental health, cultural use, and economic value. | o | o |
| 3. Describe how energy moves through trophic levels in an ecosystem, including the ways in which organisms use energy. | o | o |
| 4. Provide examples of biological magnification. | o | o |

| | | |
|---|---|---|
| 5. Identify the human health consequences of environmental degradation such as dioxin, water borne diseases, and poor air quality. | o | o |
| 6. Construct an argument that relates a human health issue with the environment (e.g., water-borne disease, food poisoning, air quality, dioxin, UV radiation). | o | o |

Q5.6 Thinking about the socioscientific issue of ECOSYSTEM HEALTH, is there anything students need to learn about it that is missing in this list of LOs?

o Yes

o No

Q5.7 Please explain what LOs are missing from the socioscientific issue of ECOSYSTEM HEALTH.

**Q5.8 Please indicate whether the following LOs are critical or not critical for teaching this issue to non-majors.**

| TOPIC: BIOREMEDIATION & SOIL | CRITICAL | NOT CRITICAL |
|---|---|---|
| 1. Explain the role that soil plays in terrestrial ecosystems. | o | o |
| 2. Distinguish between organic and inorganic components of soil. | o | o |
| 3. Explain why high levels of organic matter are important in soil. | o | o |
| 4. Using a list of the components of a soil, predict how well it would retain water. | o | o |
| 5. Identify environmental consequences of agricultural practices such as tilling and fertilizer use. | o | o |
| 6. Provide three benefits that mycorrhiza confer to plants. | o | o |
| 7. Compare features of soil fungi and bacteria that could make them harmful or beneficial to plants. | o | o |
| 8. Define characteristics of soils that suppress disease in plants. | o | o |
| 9. Describe the process of bioremediation (how plants can remove pollutants from the environment). | o | o |

| | | |
|---|---|---|
| 10. Design an experiment to test a question of interest related to soil restoration including choosing controls, interpreting findings, and explaining why decisions were made. | o | o |
| 11. Explain the biological process of composting. | o | o |
| 12. Given what you know about developing and maintaining a compost pile, diagnose problems with a compost pile. | o | o |
| 13. Describe the benefits of composting as a carbon sink, for agriculture, and for reducing waste. | o | o |
| 14. Be able to calculate the C:N ratios of composting materials given tables of compost rates. | o | o |
| 15. Provide examples of how compost piles can be used to produce energy. | o | o |

Q5.9 Thinking about the socioscientific issue of BIOREMEDIATION & SOIL, is there anything students need to learn about it that is missing in these list of LOs?
o Yes
o No
Q5.10 Please explain what LOs are missing from the socioscientific issue of BIOREMEDIATION & SOIL.
Q5.11 Please indicate whether the following LOs are critical or non critical for teaching this issue to non-majors.

| TOPIC: SUSTAINABLE FOOD & GMOs | CRITICAL | NOT CRITICAL |
|---|---|---|
| 1. Describe why someone would be concerned about or support genetic modifications in organisms like crops. | o | o |
| 2. Discuss the costs and benefits associated with genetically modified organisms, including human gene therapy. | o | o |
| 3. Identify two to three examples of GMO foods and the modifications introduced to create each. | o | o |
| 4. Compare two common methods used to generate genetic modifications in plants. | o | o |

| | | |
|---|---|---|
| 5. Distinguish between selective breeding and genetic modifications. | o | o |
| 6. Find and evaluate research studies that serve as the basis for understanding the safety of genetic manipulations in crops, and use these scientific resources to support an argument or decision. | o | o |

Q5.12 Thinking about the socioscientific issue of SUSTAINABLE FOOD & GMOs, is there anything students need to learn about it that is missing in this list of LOs?
o Yes
o No
Q5.13 Please explain what LOs are missing from the socioscientific issue of SUSTAINABLE FOOD & GMOs.
**End of Block: Ecology (Biodiversity loss, soil, GMOs, sustainable food)**

**Start of Block: Ecology (Overfishing, Eutrophication, Ocean acidification, plastic pollution)**
Q6.1 This block of 33 learning objectives is part of the Ecology unit and contain the issues:
OVERFISHING
EUTROPHICATION
CORAL BLEACHING
OCEAN ACIDIFICATION & PLASTIC POLLUTION
Please note that the learning objectives you are evaluating are for this a socioscientific issue. Consider if they are important for students to learn and if these LOs encompass what students need to learn about this issue.

Q6.2 Please indicate whether the following LOs are critical or not critical for teaching this issue to non-majors.
TOPIC: OVERFISHING

| | CRITICAL | NOT CRITICAL |
|---|---|---|
| 1. Identify abiotic factors that influence aquatic biomes. | o | o |
| 2. Compare the characteristics of ocean zones and identify major types of organisms that would be present in different zones. | o | o |
| 3. Identify trends in data (e.g., population size and density) on species in a habitat. | o | o |
| 4. Analyze the role of organisms as producers, consumers, and decomposers in marine ecosystems. | o | o |

| | | |
|---|---|---|
| 5. Develop and use a model of the movement of energy in marine ecosystems to evaluate the impact of an environmental perturbation. | o | o |
| 6. Justify how water temperature, depth, stratum, location, and/or time of year affect the abundance of organisms at a location. | o | o |
| 7. Predict and explain your reasoning about where you would find a specific species in a particular habitat, if given information on the natural history of a species. | o | o |
| 8. Compare different perspectives of stakeholders involved in regulating fisheries. | o | o |

Q6.3 Thinking about the socioscientific issue of OVERFISHING, is there anything students need to learn about it that is missing in this list of LOs?
o Yes
o No

Q6.4 Please explain what LOs are missing from the socioscientific issue of OVERFISHING.

Q6.5 Please indicate whether the following LOs are critical or not critical for teaching this issue to non-majors.

| TOPIC: EUTROPHICATION | CRITICAL | NOT CRITICAL |
|---|---|---|
| 1. Provide an example of why nitrogen is needed for living organisms. | o | o |
| 2. Compare the diverse ways that living organisms acquire nitrogen. | o | o |
| 3. Construct a model of the nitrogen cycle in an ecosystem. | o | o |
| 4. Define “dead zones” and eutrophication. | o | o |
| 5. Identify human behaviors that lead to excess nitrogen in the environment. | o | o |
| 6. Predict how increasing nitrogen impacts microbes in an ecosystem. | o | o |
| 7. Track how nutrients in human waste could be reclaimed. | o | o |
| 8. Critique approaches to human waste in terms of environmental sustainability. | o | o |

Q6.6 Thinking about the socioscientific issue of EUTROPHICATION, is there anything students need to learn about it that is missing in this list of LOs?
o Yes
o No

Q6.7 Please explain what LOs are missing from the socioscientific issue of EUTROPHICATION.

Q6.8 Please indicate whether the following LOs are critical or not critical for teaching this issue to non-majors.

| TOPIC: CORAL BLEACHING | **CRITICAL** | **NOT CRITICAL** |
|---|---|---|
| 1. Explain the benefits of the symbiotic relationship between corals and zooxanthellae. | **o** | **o** |
| 2. Characterize the locations of coral reef ecosystems around the world. | **o** | **o** |
| 3. Explain the consequences of rising sea surface temperature on coral reefs. | **o** | **o** |
| 4. Interpret sea surface temperature maps to understand the range and habit of coral reefs. | **o** | **o** |
| 5. Calculate accumulated heat stress from sea surface temperature data. | **o** | **o** |
| 6. Graph and interpret data to estimate potential harm to an ecosystem. | **o** | **o** |
| 7. Analyze a graph to predict the occurrence, timing and severity of coral bleaching. | **o** | **o** |

Q6.9 Thinking about the socioscientific issue of CORAL BLEACHING, is there anything students need to learn about it that is missing in this list of LOs?
o Yes
o No

Q6.10 Please explain what LOs are missing from the socioscientific issue of CORAL BLEACHING.

Q6.11 Please indicate whether the following LOs are critical or not critical for teaching this issue to non-majors.

| TOPIC: OCEAN ACIDIFICATION & PLASTICS | CRITICAL | NOT CRITICAL |
|---|---|---|
| 1. Read and interpret graphs of atmospheric and ocean $CO_2$. | o | o |

| | | |
|---|---|---|
| 2. Explain the inputs, outputs, and sources of chemical reactions that occur with CO2 in seawater that affect available calcium carbonate for shell-building organisms | o | o |
| 3. Predict the likely effects of changes of increased CO2 on ocean pH. | o | o |
| 4. Analyze ocean chemistry data to compare coastal and ocean acidification. | o | o |
| 5. Given a research question about ocean acidification, gather data to answer it. | o | o |
| 6. Describe the environmental challenges of plastic pollution. | o | o |
| 7. Interpret graphs and maps of plastic pollution. | o | o |
| 8. Characterize how plastic and other materials should be appropriately recycled. | o | o |
| 9. Evaluate proposed models of plastic pollution reduction. | o | o |
| 10. Explain characteristics of microbes that can decompose plastic. | o | o |

Q6.12 Thinking about the socioscientific issue of OCEAN ACIDIFICATION & PLASTICS, is there anything students need to learn about it that is missing in this list of LOs?
o Yes
o No
Q6.13 Please explain what LOs are missing from the socioscientific issue of OCEAN ACIDIFICATION & PLASTICSn.
**End of Block: Ecology (Overfishing, Eutrophication, Ocean acidification, plastic pollution)**

**Start of Block: Evolution (Antibiotic Resistance, Microbiomes)**
Q7.1 This block of 22 learning objectives is part of the Evolution unit and contain the issues:
ANTIBIOTIC RESISTANCE
MICROBIOMES
Please note that the learning objectives you are evaluating are for this a socioscientific issue. Consider if they are important for students to learn and if these LOs encompass what students need to learn about this issue.
Q7.2 Please indicate whether the following LOs are critical or not critical for teaching this issue to non-majors.

| TOPIC: ANTIBIOTIC RESISTANCE | CRITICAL | NOT CRITICAL |
|---|---|---|
| 1. Given a description of a cell or virus, be able to predict features that are sensitive to antibiotics. | o | o |
| 2. Diagram the events of the evolution of antibiotic resistance with a focus on mutation, selective advantage, reproductive advantage, and population level outcomes. | o | o |
| 3. Identify different ways that bacteria can pass new genetic material on to other bacteria. | o | o |
| 4. Compare vertical vs. horizontal genetic transfer of information. | o | o |
| 5. Given information on where an antibiotic binds to molecules essential for transcription or translation, explain 1) why the antibiotic disrupts the process, and 2) why the disruption affects only bacterial cells--not eukaryotic cells. | o | o |
| 6. Analyze the protein produced from different alleles to support a claim of antibiotic resistance or susceptibility. | o | o |
| 7. Provide an example of evolution in a population. | o | o |
| 8. Distinguish between the idea that selection acts on individuals but only populations evolve. | o | o |
| 9. List the critical steps for evolution to occur by natural selection. | o | o |
| 10. Explain how DNA sequences can be used to trace the evolution and spread of antibiotic resistance. | o | o |
| 11. Define phylogeny and be able to infer relationships from reading a phylogeny. | o | o |
| 12. Use phylogenies to make conclusions about the relationships between isolates of a bacterium. | o | o |
| 13. Provide examples of antibiotic-resistant bacteria and explain why antibiotic resistance is a problem. | o | o |

Q7.3 Thinking about the socioscientific issue of ANTIBIOTIC RESISTANCE, is there anything students need to learn about it that is missing in this list of LOs?
o Yes

o No

Q7.4 Please explain what LOs are missing from the socioscientific issue of ANTIBIOTIC RESISTANCE.

Q7.5 Please indicate whether the following LOs are critical or not critical for teaching this issue to non-majors.

| TOPIC: MICROBIOMES | CRITICAL | NOT CRITICAL |
|---|---|---|
| 1. Compare the size, structure, mode of reproduction, genetic material, and other features of viruses to prokaryotic and eukaryotic cells. | o | o |
| 2. Compare and contrast key elements of bacterial versus eukaryotic cell structure. | o | o |
| 3. Distinguish the features of prokaryotes that differ from eukaryotes or viruses. | o | o |
| 4. Consider DNA replication, transcription, and translation, and for each process explain at least one key difference and one similarity in how they occur in bacterial versus eukaryotic cells. | o | o |
| 5. Explain the role of microbiomes in human health. | o | o |
| 6. Differentiate between mutualism, commensalism, and parasitism and recognize examples of these in human-bacterial symbiotic relationships. | o | o |
| 7. Explain the difference between species diversity and richness and recognize examples of each. | o | o |
| 8. Identify organisms of concern based on data from hospitalizations and deaths from foodborne illnesses. | o | o |
| 9. Explain the techniques of PCR and sequence alignment that the CDC uses to determine the pathogenic organism. | o | o |

Q7.6 Thinking about the socioscientific issue of MICROBIOMES, is there anything students need to learn about it that is missing in this list of LOs?

o Yes

o No

Q7.7 Please explain what LOs are missing from the socioscientific issue of MICROBIOMES.

**End of Block: Evolution (Antibiotic Resistance, Microbiomes)**

**Start of Block: Genetics (Inheritance & Genetic Testing)**
Q8.1 This block of 48 learning objectives (LOs) are part of the Genetics unit and contain the issues:
INHERITANCE
GENETIC TESTING
NON-BIOLOGICAL BASIS OF RACE
You are evaluating the LOs for these socioscientific issues. Consider if the LOs are critical for students to learn and if these LOs encompass what students need to learn about this issue.
Q8.2 Please indicate whether the following LOs are critical or not critical for teaching this issue to non-majors.

| TOPIC: INHERITANCE | CRITICAL | NOT CRITICAL |
|---|---|---|
| 1. Define phenotype, genotype, dominant, recessive, heterozygous, and homozygous. | o | o |
| 2. Compare the genetic information held on two homologous chromosomes, two nonhomologous chromosomes and two sister chromatids. | o | o |
| 3. Define the following terms: CHROMOSOME--and distinguish replicated and unreplicated--CHROMATID, PLOIDY, HAPLOID NUMBER, HOMOLOGOUS CHROMOSOMES. | o | o |
| 4. Given a micrograph or drawing of a cell you've never seen before, label chromosomes, chromatids, sister chromatids, and homologous chromosomes, if present, and determine the haploid number and ploidy of the cell. | o | o |
| 5. Explain how the combination of alleles determines phenotype for a simple monogenic trait with a dominant/recessive pattern of inheritance. | o | o |
| 6.Calculate the probability of a particular gamete being produced from an individual, assuming independent segregation. | o | o |

| | | |
|---|---|---|
| 7. Calculate the probability of a particular genotype, given independent segregation and random union of gametes between two individuals with known genotypes. | o | o |
| 8. Label which elements in a Punnett square represent the genotypes of egg, sperm, and offspring. Explain how you can determine the frequency of each egg and sperm genotype for one trait and how you can use this information to calculate the frequencies of offspring genotypes and phenotypes. | o | o |
| 9. Given any pair of parental genotypes and information on the alleles present, use a Punnett square to complete a genetic cross. Identify the genotypes and phenotypes of offspring and calculate their predicted frequencies. Note that the genes involved may be autosomal, X-linked, linked, or unlinked and that the alleles involved may be dominant, recessive, or co-dominant. | o | o |
| 10. Given information about two parents, use Punnett squares to determine the probability that a child will inherit (1) an autosomal dominant, (2) an autosomal recessive, or (3) a sex-linked recessive allele and/or show symptoms of the disease caused by this allele if given information about the parents. | o | o |
| 11. Given a completed Punnett square and information on offspring phenotypes, determine whether the alleles involved are 1) dominant, recessive, or codominant, 2) autosomal or X-linked, and 3) linked or unlinked. | o | o |
| 12. On a pedigree, label 1) males and females, 2) affected and unaffected individuals, and 3) generations. | o | o |
| 13. Interpret pedigree information to determine the suitability of a DNA marker for tracking a disease trait in a family. | o | o |
| 14. Using pedigrees, distinguish between dominant, recessive, autosomal, X-linked monogenic traits. | o | o |
| 15. Calculate the probability that an individual in a pedigree has a particular genotype. | o | o |

Q8.3 Thinking about the socioscientific issue of INHERITANCE, is there anything students need to learn about it that is missing in these list of LOs?
o Yes
o No
Q8.4 Please explain what LOs are missing from the socioscientific issue of INHERITANCE.

Q8.5 Please indicate whether the following LOs are critical or non critical for teaching this issue to non-majors.

| GENETIC TESTING | CRITICAL | NOT CRITICAL |
|---|---|---|
| 1. Describe different types of genetic variations (mutations) in the genome, including their location in coding and noncoding DNA. | o | o |
| 2. If given information about the type and genome location of an inherited genetic variation, predict whether it will alter the structure or amount of a protein inside cells, and whether the genetic variation is likely to change a trait. | o | o |
| 3. Rank the following mutations in terms of greatest to least impact on the structure and function of genes and gene products: missense (change amino acids), nonsense (change to "stop"), frameshift (change reading frame), and silent (no change in product). Explain your reasoning. | o | o |
| 4. Predict how a given mutation in DNA may or may not affect that amino acid sequence of the protein produced (through transcription and translation.) | o | o |
| 5. Explain the difference between chromosomes, genes, DNA, RNA, and proteins. | o | o |
| 6. Defend the statements: "The information in DNA is heritable. The information in proteins is not." | o | o |
| 7. Explain how continuous traits involve many different genes, each with alleles that can contribute varying amounts to the phenotype. | o | o |
| 8. Evaluate how genes and the environment can interact to produce a phenotype. | o | o |

| | | |
|---|---|---|
| 9. Describe how scientists use Genome-Wide Association Studies (GWAS) to determine how certain gene variants contribute to complex diseases. | o | o |
| 10. Given information about a gene variant (allele) and data from GWAS, calculate the likelihood of genetic predispositions. | o | o |
| 11.Explain the purpose of a genetic test. | o | o |
| 12. Discuss the ethical issues involved in obtaining and using data on DNA sequences and chromosome structure in human parents and fetuses. | o | o |
| 13. Research real-life genetic testing scenarios and consider the advantages and disadvantages associated with common genetic tests for different traits. | o | o |
| 14. Explain the role and stepwise process in using PCR, gel electrophoresis, SNP chips, and gene sequencing in documenting genotypes. | o | o |
| 15. Choose a test to use based on what is known about a genetic trait or condition and interpret its results. | o | o |
| 16. Identify genetic and environmental factors linked to the development of a disorder. | o | o |
| 17. Research genetic and environmental factors linked to the development of a human trait using materials on direct-to-consumer genetic testing sites. | o | o |
| 18. Describe a study design that examined a link with a human trait and a gene. | o | o |
| 19. Given a data table, identify the estimated increase in propensity to develop a human trait if someone were to have a mutation in a specific gene. | o | o |
| 20. Describe the product from a gene linked to a human trait and explain why some alleles are associated with different reactions. | o | o |

Q8.6 Thinking about the socioscientific issue of GENETIC TESTING, is there anything students need to learn about it that is missing in this list of LOs?

o Yes

o No

Q8.7 Please explain what LOs are missing from the socioscientific issue of GENETIC TESTING.

Q8.8 Please indicate whether the following LOs are critical or non critical for teaching this issue to non-majors.

| NON-BIOLOGICAL BASIS OF RACE | CRITICAL | NOT CRITICAL |
|---|---|---|
| 1. Provide examples of the social history of race. | o | o |
| 2. Use evidence to defend the statement that race cannot be defined using biological traits. | o | o |
| 3. Compare the biological definition of genomic ancestry and the social definition of race. | o | o |
| 4. Use evidence to explain why light or dark skin color is beneficial or harmful in a particular environment, especially before vitamin D supplements. | o | o |
| 5. Use what you know about UV radiation and DNA damage, to discuss the role of melanin, mutations and skin cancer. | o | o |
| 6. Analyze examples where race is useful and important in biomedical research—and how it is not useful. | o | o |
| 7. Explain the causes and consequences of a racial health disparity. | o | o |
| 8. Make an evidence-based recommendation to reduce racial health disparities. | o | o |
| 9. Infer an individual's ancestry given information on alleles associated with skin color or other traits. | o | o |
| 10. Use allele frequencies to infer ancestry (e.g. skin color-related alleles). | o | o |
| 11. Interpret DNA evidence to explain patterns of genetic variation in human ancestry and migration out of Africa. | o | o |
| 12. Using a map of the world, explain the Out of Africa hypothesis, including the role of genetic drift, natural selection, and gene flow in the evolution of Homo sapiens. | o | o |
| 13. Critique the use of physical traits to define race. | o | o |

Q8.9 Thinking about the socioscientific issue of NON-BIOLOGICAL BASIS OF RACE, is there anything students need to learn about it that is missing in this list of LOs?
o Yes
o No

Q8.10 Please explain what LOs are missing from the socioscientific issue of NON-BIOLOGICAL BASIS OF RACE

**End of Block: Genetics (Inheritance & Genetic Testing)**

**Start of Block: Physiology (HIV, Pandemics, Vaccines, and Allergies)**

Q9.1 This block of 25 learning objectives is part of the Physiology (Immune System) unit and contain the issues:
HIV & VIRUSES
PANDEMICS & VACCINES
ALLERGIES
Please note that the learning objectives you are evaluating are for this a socioscientific issue. Consider if they are important for students to learn and if these LOs encompass what students need to learn about this issue.

Q9.2 Please indicate whether the following LOs are critical or not critical for teaching this issue to non-majors.

| TOPIC: HIV & VIRUSES | CRITICAL | NOT CRITICAL |
|---|---|---|
| 1. Describe a common sexually transmitted disease: how it is transmitted, its characteristics, symptoms, and rate of spread. | o | o |
| 2. Evaluate misconceptions about how HIV/AIDs is transmitted. | o | o |
| 3. Identify the functions of major proteins of a virion. | o | o |
| 4. Explain how viral receptors and cell surface proteins modulate the specificity of viral cell entry and infection. | o | o |
| 5. Define viral hosts and vectors. | o | o |
| 6. Analyze viral genetic sequences across space and time to make inferences about disease transmission over time. | o | o |
| 7. Define the function of these major cell types in the immune system: B cells, cytotoxic T cells, helper T cells, antigen presenting cells, macrophages, natural killer cells. | o | o |

Q9.3 Thinking about the socioscientific issue of HIV & VIRUSES, is there anything students need to learn about it that is missing in this list of LOs?
o Yes (1)
o No (2)

Q9.4 Please explain what LOs are missing from the socioscientific issue of HIV & VIRUSES.
Q9.5 Please indicate whether the following LOs are critical or not critical for teaching this issue to non-majors.

| TOPIC: PANDEMICS & VACCINES | CRITICAL | NOT CRITICAL |
|---|---|---|
| 1. Define pandemic, case fatality rate, incubation period. | o | o |
| 2. Interpret data about viral stability, case fatality rate, number of people who remain asymptomatic for an infection to make predictions about disease outcomes. | o | o |
| 3. Distinguish between primary and secondary immune responses. | o | o |
| 4. Define antibodies and explain how they target antigens for destruction. | o | o |
| 5. Define herd immunity. | o | o |
| 6. Define the reproduction number, R0, of a virus. | o | o |
| 7. Explain how herd immunity relates to vaccination and R0. | o | o |
| 8. Evaluate reproduction rates and vaccination rates needed to reach herd immunity. | o | o |
| 9. Describe viral vector, live attenuated, and inactivated vaccines. | o | o |
| 10. Compare the advantages and disadvantages associated with certain vaccines. | o | o |
| 11. Evaluate features of a vaccine that provide value to society. | o | o |
| 12. Describe the role of random sampling, double-blinding, and control treatments in the design of a clinical trial for a vaccine. | o | o |

| | | |
|---|---|---|
| 13. Critique methodological and ethical issues with the Wakefield Lancet study. | o | o |
| 14. Debate the ethics of COVID vaccine development and distributions. | o | o |

Q9.6 Thinking about the socioscientific issue of PANDEMICS & VACCINES, is there anything students need to learn about it that is missing in this list of LOs?
o Yes
o No
Q9.7 Please explain what LOs are missing from the socioscientific issue of PANDEMICS & VACCINES.

Q9.8 Please indicate whether the following LOs are critical or not critical for teaching this issue to non-majors.

| TOPIC: ALLERGIES | CRITICAL | NOT CRITICAL |
|---|---|---|
| 1. Describe the role of mast cells, histamines, and IgE antibodies in an allergic response. | o | o |
| 2. Compare an allergic response, including mast cells, histamine, IgE antibodies, with a normal immune response. | o | o |
| 3. Describe inferences and conclusions from allergen study data. | o | o |
| 4. Define RNA interference and describe how it can be used to affect gene function. | o | o |

Q9.9 Thinking about the socioscientific issue of ALLERGIES, is there anything students need to learn about it that is missing in this list of LOs?
o Yes
o No

Q9.10 Please explain what LOs are missing from the socioscientific issue of ALLERGIES.
**End of Block: Physiology (HIV, Pandemics, Vaccines, and Allergies)**

**Start of Block: Physiology (Nervous System, Stress, Anxiety, and Depression)**
Q10.1 This block of 18 learning objectives is part of the Physiology (Nervous System) unit and contain the issues:
NERVOUS SYSTEM
ANXIETY & DEPRESSION
Please note that the learning objectives you are evaluating are for this a socioscientific issue.

Consider if they are important for students to learn and if these LOs encompass what students need to learn about this issue.

Q10.2 Please indicate whether the following LOs are critical or not critical for teaching this issue to non-majors.

TOPIC: NERVOUS SYSTEM

| | CRITICAL | NOT CRITICAL |
|---|---|---|
| 1. Explain why ions and polar molecules do not move across plasma membranes efficiently. | o | o |
| 2. Predict the relative rates at which given ions and molecules will cross a plasma membrane in the absence of membrane proteins. Explain your reasoning. | o | o |
| 3. Explain how an electrochemical gradient is maintained across a plasma membrane. | o | o |
| 4. Predict 1) which direction a given ion or molecule will cross a plasma membrane given an electrochemical gradient, and 2) whether a channel, carrier, or pump would be involved. | o | o |
| 5. Describe the parts of a neuron including dendrite, axon, cell body, synaptic cleft, post-synaptic cleft, myelin. | o | o |
| 6. Describe how neurons communicate with each other using neurotransmitters. | o | o |
| 7. Identify specific neurotransmitters and their effects in the body. | o | o |
| 8. Describe the roles of the two parts of the autonomic nervous system. | o | o |
| 9. Identify the functions of motor neurons, interneurons, and sensory neurons. | o | o |
| 10. Given a scenario, explain how the central nervous system and peripheral nervous system process and respond to sensory information. | o | o |
| 11. Describe how a stimulus triggers a sensory cell to send a message electrochemically. | o | o |

Q10.3 Thinking about the socioscientific issue of the NERVOUS SYSTEM, is there anything students need to learn about it that is missing in this list of LOs?
o Yes
o No
Q10.4 Please explain what LOs are missing from the socioscientific issue of NERVOUS SYSTEM.

Q10.5 Please indicate whether the following LOs are critical or not critical for teaching this issue to non-majors.

| TOPIC: ANXIETY & DEPRESSION | CRITICAL | NOT CRITICAL |
|---|---|---|
| 1. Explain what happens to the body while it is in fight, flight, or freeze. | o | o |
| 2. Explain how variation in heart rate is used as an index for stress. | o | o |
| 3. Given a specific recreational or abused drug, describe its impacts on the nervous system. | o | o |
| 4. Describe biologically how a stress cycle can be completed. | o | o |
| 5. Examine the efficacy of stress-relief strategies designed to stimulate the parasympathetic nervous system and calm the sympathetic nervous system. | o | o |
| 6. Describe the characteristics of the key neurotransmitters implicated in depression. | o | o |
| 7. Examine scientific studies to determine the efficacy for treatments for depression. | o | o |

Q10.6 Thinking about the socioscientific issue of ANXIETY & DEPRESSION, is there anything students need to learn about it that is missing in this list of LOs?
o Yes
o No

Q10.7 Please explain what LOs are missing from the socioscientific issue of ANXIETY & DEPRESSION.
**End of Block: Physiology (Nervous System, Stress, Anxiety, and Depression)**

**Start of Block: Physiology (Human sensory experiences, disability)**
Q11.1 This block of 17 learning objectives is part of the Physiology (Sensory experiences & Disability) unit and contain the issues:
PAIN
DISABILITY (HEARING, VISION, TASTE)
Please note that the learning objectives you are evaluating are for this a socioscientific issue.

Consider if they are important for students to learn and if these LOs encompass what students need to learn about this issue.

Q11.2 Please indicate whether the following LOs are critical or not critical for teaching this issue to non-majors.

| TOPIC: PAIN | CRITICAL | NOT CRITICAL |
|---|---|---|
| 1. Compare variation in sensory experiences of non-human organisms. | o | o |
| 2. Explore what we know about variation in nociception in humans. | o | o |
| 3. Explain why variation in pain is important for medical care. | o | o |
| 4. Describe how opioids work. | o | o |
| 5. Describe how mechanoreceptors work. | o | o |
| 6. Create a diagram to predict how space travel impacts mechanoreceptors in bone remodeling and calcium regulation, including the effects on bone health. | o | o |

Q11.3 Thinking about the socioscientific issue of PAIN, is there anything students need to learn about it that is missing in this list of LOs?

o Yes

o No

Q11.4 Please explain what LOs are missing from the socioscientific issue of PAIN.

Q11.5 Please indicate whether the following LOs are critical or not critical for teaching this issue to non-majors.

| TOPIC: DISABILITY (HEARING, VISION, TASTE) | CRITICAL | NOT CRITICAL |
|---|---|---|
| 1. Identify the structures of the auditory system and their functions. | o | o |
| 2. Identify the structures of the vestibular system and their functions. | o | o |
| 3. Identify the structures involved in vision and their functions. | o | o |

| | | |
|---|---|---|
| 4. Identify the structures and functions involved in taste. | o | o |
| 5. Predict the evolutionary advantage of tasting bitter substances. | o | o |
| 6. Identify the structures and functions involved in taste. | o | o |
| 7. Predict the evolutionary advantage of tasting bitter substances. | o | o |
| 8. Provide an example of a gene that affects taste. | o | o |
| 9. Identify the structures and functions involved in smell. | o | o |
| 10. Generate hypotheses that could explain why smell is important to humans. | o | o |
| 11. Provide an example of why neuroplasticity is important to human functioning. | o | o |

Q11.6 Thinking about the socioscientific issue of DISABILITY (HEARING, VISION, TASTE), is there anything students need to learn about it that is missing in this list of LOs?
o Yes
o No
Q11.7 Please explain what LOs are missing from the socioscientific issue of DISABILITY (HEARING, VISION, TASTE)
**End of Block: Physiology (Human sensory experiences, disability)**

**Start of Block: Physiology (Sex, reproduction, & gender)**
Q12.1 This block of 12 learning objectives is part of the Physiology (Sex, reproduction, & gender) unit and contain the issues:
GENDER
FERTILITY
Please note that the learning objectives you are evaluating are for this a socioscientific issue. Consider if they are important for students to learn and if these LOs encompass what students need to learn about this issue.
Q12.2 Please indicate whether the following LOs are critical or not critical for teaching this issue to non-majors.

| TOPIC: GENDER | CRITICAL | NOT CRITICAL |
|---|---|---|
| 1. Describe the role of sex chromosomes to determine biological sex in humans. | o | o |

| | | |
|---|---|---|
| 2. Evaluate the ethical, personal, and societal implications of sex verification tests. | o | o |
| 3. Describe variation in sexual orientation in humans. | o | o |
| 4. Distinguish between sex, gender, and sexual orientation. | o | o |
| 5. Describe variation in sex, gender, and sexuality in organisms beyond humans. | o | o |
| 6. Generate a hypothesis about sex, gender, or sexuality in a non-human organism. | o | o |

Q12.3 Thinking about the socioscientific issue of GENDER, is there anything students need to learn about it that is missing in this list of LOs?
o Yes (1)
o No (2)
Q12.4 Please explain what LOs are missing from the socioscientific issue of GENDER.

Q12.5 Please indicate whether the following LOs are critical or not critical for teaching this issue to non-majors.

| TOPIC: FERTILITY | CRITICAL | NOT CRITICAL |
|---|---|---|
| 1. Identify how, where, and when sperm and eggs are produced. | o | o |
| 2. Compare and contrast the ovarian and uterine cycle. | o | o |
| 3. Describe the major structures and functions of the reproductive systems. | o | o |
| 4. Identify the role of estrogen, progesterone, and testosterone in human reproductive systems. | o | o |
| 5. Given change to estrogen, progesterone, and testosterone levels, predict the outcome for human reproduction, for example, to manipulate fertility or for gender confirmation. | o | o |
| 6. Given a calendar of a person's menstrual cycle, use your knowledge of the ovarian and uterine cycles, ovulation, and fertilization to predict when ovulation and pregnancy are most likely. | o | o |

Q12.6 Thinking about the socioscientific issue of FERTILITY, is there anything students need to learn about it that is missing in this list of LOs?

o Yes
o No = ***Yes***

Q12.7 Please explain what LOs are missing from the socioscientific issue of FERTILITY
**End of Block: Physiology (Sex, reproduction, & gender)**

Q13.6 If you have any questions about the project or this survey, please do not hesitate to contact us.

### Round 3 - Asynchronous Collaborative Discussion of Survey Results

#### Email:

Title: Final round of review for a set of learning objectives for non-majors' biology

Dear ___,
Thank you for your willingness to help evaluate and curate a nascent list of learning objectives for non-majors' biology. To aid us in the final step of our validation, please help us one last time by clicking on the files below to view:

1. A list of the LOs that were rated less critical for the topic indicated. If you have any additional comments to make on these orphan LOs. **(1) indicate if you think the concept should be included in our final set of official LOs for non-majors' biology and (2) explain your reasoning using the comment feature**
2. A list of the additions that this community suggested. **(1) indicate if you think the concept should be included in our final set of official LOs for non-majors' biology and (2) explain your reasoning using the comment feature.**
3. A list of the LOs deemed most critical to teaching these topics to non-majors. Thank you and enjoy reading the level of consensus.

Please comment in the next two weeks. This last step will hopefully help us get these published more widely for the whole community to access.

##### Sample Google Doc for Collecting Comments

Task #1:
**Reviewers struggled to determine if following LOs were critical or not. For each LO, please use the comment function: (1) indicate if you think the LO should be included in our final set of official LOs for non-major's biology and (2) explain your reasoning.**
Green - Lower Level Blooms items
Blue - Higher Level Blooms items
**For your reference, the following LOs have >70% agreement - Critical. There is no need to comment on these LOs.**

### Genetics (Inheritance & Genetic Testing)

#### Task #1:

**Reviewers struggled to determine if following LOs were critical or not. For each LO, please use the comment function: (1) indicate if you think the LO should be included in our final set of official LOs for non-major's biology and (2) explain your reasoning.**

Green - Lower Level Blooms items

Blue - Higher Level Blooms items

| INHERITANCE | Critical/ Not Critical | |
|---|---|---|
| On a pedigree, label 1) males and females, 2) affected and unaffected individuals, and 3) generations. | 4 | 4 |
| Given a micrograph or drawing of a cell you've never seen before, label chromosomes, chromatids, sister chromatids, and homologous chromosomes, if present, and determine the haploid number and ploidy of the cell. | 2 | 6 |
| Interpret pedigree information to determine the suitability of a DNA marker for tracking a disease trait in a family. | 3 | 5 |
| Calculate the probability that an individual in a pedigree has a particular genotype. | 3 | 5 |
| GENETIC VARIATIONS | | |
| Describe how scientists use Genome-Wide Association Studies (GWAS) to determine how certain gene variants contribute to complex diseases. | 5 | 3 |
| Defend the statements: "The information in DNA is heritable. The information in proteins is not." | 4 | 4 |
| Choose a test to use based on what is known about a genetic trait or condition and interpret its results. | 4 | 4 |
| Research genetic and environmental factors linked to the development of a human trait using materials on direct-to-consumer genetic testing sites. | 4 | 4 |
| Given a data table, identify the estimated increase in propensity to develop a human trait if someone were to have a mutation in a specific gene. | 4 | 4 |
| Explain the role and stepwise process in using PCR, gel electrophoresis, SNP chips, and gene sequencing in documenting genotypes. | 3 | 5 |

| | |
|---|---|
| Given information about a gene variant (allele) and data from GWAS, calculate the likelihood of genetic predispositions. | 2 / 6 |
| NON-BIOLOGICAL BASIS OF RACE | Critical/ Not Critical |
| Make an evidence-based recommendation to reduce racial health disparities. | 4 / 2 |
| Interpret DNA evidence to explain patterns of genetic variation in human ancestry and migration out of Africa. | 4 / 2 |
| Using a map of the world, explain the Out of Africa hypothesis, including the role of genetic drift, natural selection, and gene flow in the evolution of Homo sapiens. | 4 / 2 |
| Infer an individual's ancestry given information on alleles associated with skin color or other traits. | 3 / 3 |
| Use allele frequencies to infer ancestry (e.g. skin color-related alleles). | 3 / 3 |

#### Task #2:

**Reviewers recommended adding LOs about the following content. For each concept, please use the comment function: (1) indicate if you think the concept should be included in our final set of official LOs for non-major's biology and (2) explain your reasoning.**

| INHERITANCE |
|---|
| Distinguish between the biological determinism versus biological potential |
| Define incomplete dominance, pleiotropy, epistasis, expressivity, penetrance, and the use of wildtype to indicate phenotypic or allelic frequency. |

**For your reference, the following LOs have >70% agreement - Critical. There is no need to comment on these LOs.**

Green - Lower Level Blooms

Blue - Higher Level Blooms

| INHERITANCE | Critical/ Not Critical |
|---|---|
| Define phenotype, genotype, dominant, recessive, heterozygous, and homozygous. | 8 |
| Compare the genetic information held on two homologous chromosomes, two nonhomologous chromosomes and two sister chromatids. | 7 1 |
| Explain how the combination of alleles determines phenotype for a simple monogenic trait with a dominant/recessive pattern of inheritance | 7 1 |
| Calculate the probability of a particular gamete being produced from an individual, assuming independent segregation. | 7 1 |
| Calculate the probability of a particular genotype, given independent segregation and random union of gametes between two individuals with known genotypes. | 7 1 |
| Label which elements in a Punnett square represent the genotypes of egg, sperm, and offspring. Explain how you can determine the frequency of each egg and sperm genotype for one trait and how you can use this information to calculate the frequencies of offspring genotypes and phenotypes. | 7 1 |
| Given any pair of parental genotypes and information on the alleles present, use a Punnett square to complete a genetic cross. Identify the genotypes and phenotypes of offspring and calculate their predicted frequencies. Note that the genes involved may be autosomal, X-linked, linked, or unlinked and that the alleles involved may be dominant, recessive, or co-dominant. | 7 1 |
| Given information about two parents, use Punnett squares to determine the probability that a child will inherit (1) an autosomal dominant, (2) an autosomal recessive, or (3) a sex-linked recessive allele and/or show symptoms of the disease caused by this allele if given information about the parents | 7 1 |
| Given a completed Punnett square and information on offspring phenotypes, determine whether the alleles involved are 1) dominant, recessive, or codominant, 2) autosomal or X-linked, and 3) linked or unlinked. | 7 1 |
| Using pedigrees, distinguish between dominant, recessive, autosomal, X-linked monogenic traits | 6 2 |
| Define the following terms: CHROMOSOME--and distinguish replicated and unreplicated--CHROMATID, PLOIDY, HAPLOID NUMBER, HOMOLOGOUS CHROMOSOMES | 6 2 |
| GENETIC VARIATIONS | Critical/ Not Critical |
| Explain how continuous traits involve many different genes, each with alleles that can contribute varying amounts to the phenotype. | 8 |

| | |
|---|---|
| Evaluate how genes and the environment can interact to produce a phenotype. | 8 |
| Rank the following mutations in terms of greatest to least impact on the structure and function of genes and gene products: missense (change amino acids), nonsense (change to "stop"), frameshift (change reading frame), and silent (no change in product). Explain your reasoning. | 8 1 |
| Explain the difference between chromosomes, genes, DNA, RNA, and proteins | 8 1 |
| Describe different types of genetic variations (mutations) in the genome, including their location in coding and noncoding DNA. | 7 2 |
| If given information about the type and genome location of an inherited genetic variation, predict whether it will alter the structure or amount of a protein inside cells, and whether the genetic variation is likely to change a trait. | 7 2 |
| Predict how a given mutation in DNA may or may not affect that amino acid sequence of the protein produced (through transcription and translation) | 7 1 |
| Discuss the ethical issues involved in obtaining and using data on DNA sequences and chromosome structure in human parents and fetuses. | 7 1 |
| Research real-life genetic testing scenarios and consider the advantages and disadvantages associated with common genetic tests for different traits. | 7 1 |
| Identify genetic and environmental factors linked to the development of a human trait. | 6 2 |
| Describe a study design that examined a link with a human trait and a gene. | 6 2 |
| Explain the purpose of a genetic test. | 6 2 |
| Describe the product from a gene linked to a human trait and explain why some alleles are associated with different reactions. | 6 2 |
| NON-BIOLOGICAL BASIS OF RACE | Critical/ Not Critical |
| Critique the use of physical traits to define race. | 6 |
| Analyze examples where race is useful and important in biomedical research—and how it is not useful. | 6 |
| Use evidence to defend the statement that race cannot be defined using biological traits. | 6 |

| | |
|---|---|
| Use evidence to explain why light or dark skin color is beneficial or harmful in a particular environment, especially before vitamin D supplements. | 6 |
| Explain the causes and consequences of a racial health disparity. | 6 |
| Provide examples of the social history of race. | 5 1 |
| Compare the biological definition of genomic ancestry and the social definition of race. | 5 1 |
| Use what you know about UV radiation and DNA damage, to discuss the role of melanin, mutations and skin cancer. | 5 1 |
